# Investigating Frontoparietal Networks and Activation in Children with Mathematics Learning Difficulties: Cases with Different Deficit Profiles

**DOI:** 10.1101/2023.09.12.557321

**Authors:** Fengjuan Wang, Azilawati Jamaludin

**Affiliations:** National Institute of Education, Nanyang Technological University, Singapore

**Keywords:** Mathematics learning difficulties, graph theory, case-control design, symbolic/nonsymbolic addition

## Abstract

Approximately 15%-20% of school-aged children suffer from Mathematics Learning Difficulties (MLD). Most children with developmental dyscalculia (DD) or MLD also have comorbid cognitive deficits. Recent literature suggests that research should focus on uncovering the neural underpinnings of MLD across more inclusive samples, rather than limiting studies to pure cases of DD or MLD with highly stringent inclusion criteria. Therefore, this study aims to identify neural aberrancies that may be common across multiple MLD cases with different deficit profiles. Nine MLD cases and forty-five typically developing (TD) children, all around seven years old (27 boys), were recruited. Using functional near-infrared spectroscopy (fNIRS), brain data were collected during an approximate resting state and a mathematical computation task (addition). Graph theory was then applied to assess global and nodal network indicators of brain function. When comparing the network indicators and brain activation of the MLD cases to those of TD children, no unified neural aberrancy was found across all cases. However, three MLD cases did show distinct neural aberrancies compared to TD children. The study discusses the implications of these findings, considering both the neural aberrancies in the three MLD cases and the neural similarities found in the other six cases, which were comparable to those of the TD children. This raises important questions about the presence and nature of aberrant neural indicators in MLD across large cohorts and highlights the need for further research in this area.

## 1. Introduction

Mathematics Learning Difficulties (MLD) affect approximately 15%-20% of school-aged children and are characterized by significant challenges in learning math-related content (Ministry of Education, 2020; Räsänen et al., 2019). The most severe form of MLD is developmental dyscalculia (DD), a specific learning disability marked by persistent difficulties in acquiring essential math skills, which cannot be attributed to educational deprivation (American Psychiatric Association, 2013). An estimated 4% to 7% of school-aged children suffer from DD (Butterworth, 2019). Although some literature uses the terms DD and MLD interchangeably (Mazzocco, 2007; Wong & Chan, 2019), Rubinsten and Henik (2009) clearly differentiate between the two. They define DD as a deficit in core numerical abilities, such as difficulties in processing quantities, whereas MLD arises from multiple cognitive deficits, including challenges with working memory, visual-spatial processing, and attention.

Based on stringent definitions of DD and highly selective samples, several core deficit theories have been proposed, such as the number sense deficit and the access deficit (see Träff et al., 2017). These theories suggest that children with DD or MLD exhibit an impaired ability to rapidly and approximately distinguish between two nonsymbolic quantities without exact counting (number sense deficit) and that they struggle to connect symbols with their corresponding quantity representations (access deficit). However, in practice, most children with DD or MLD also have comorbid conditions such as dyslexia—a severe language learning difficulty (Luoni et al., 2023)—and/or exhibit poor working memory (Menon, 2016) and lower IQ (Michels et al., 2018). As a result, these theories and the rigorous sampling methods used have faced criticism in recent years for overstating the “purity” of DD or MLD. Critics argue that these approaches bias theories toward overly simplistic core-deficit accounts, failing to capture the complexity of MLD (Astle & Fletcher-Watson, 2020; Astle et al., 2022). For a comprehensive review, the following sections will explore the relevant literature on both DD and MLD, considering cases that involve pure deficits in core numerical abilities as well as those with multiple cognitive deficits.

Two of the most frequently cited behavior-cognition-brain models that explain the contributing factors of mathematical performance emphasize the role of relevant brain regions, which are situated at the most fundamental level (see reviews by Butterworth et al., 2011; Rubinsten & Henik, 2009). Generally, the frontoparietal network has been identified as crucial for mathematical skills (Iuculano & Menon, 2018; Jolles et al., 2016; Menon & Chang, 2021). Within this network, the frontal regions are associated with executive functions that support mathematical tasks, such as working memory and attention (Gilmore et al., 2018). In contrast, the parietal regions are more involved in domain-specific skills, including visual-spatial abilities and magnitude processing (Hawes et al., 2019).

Neuroimaging studies on simple addition problems have shown that children with MLD or DD exhibit different brain activation patterns in key brain regions compared to TD children. However, it is important to interpret these differences with caution, as factors such as age, diagnostic criteria, the specific arithmetic task (e.g., addition, subtraction, or multiplication), and the design of the experimental paradigm can all influence brain activation during arithmetic performance. For example, in a study by Iuculano et al. (2015), MLD children aged 7.5 to 9.6 years displayed significantly stronger activations in the bilateral prefrontal cortices and inferior and superior parietal cortices compared to TD children when engaged in a one-digit addition task. In contrast, Peters et al. (2018) found that in a one-digit subtraction task, DD children aged 9 to 12 years exhibited lower activation in the frontal and parietal areas compared to their TD counterparts.

Apart from observed activation differences from arithmetic tasks, DD children may also show abnormal resting-state functional connectivity (RSFC) in large-scale brain networks (Rapin, 2016). RSFC refers to the temporal correlation of spontaneous neural activations across brain regions without any cognitive load (Biswal et al., 1995). Jolles et al. (2016) examined the functional connectivity in children with DD. They found that DD children showed enhanced connectivity between the left and the right intraparietal sulcus (IPS), between the right IPS and the right superior frontal gyrus, and between the left IPS and the left frontal lobe. A negative relationship between hyperconnectivity of the IPS with the prefrontal and the parietal cortex and mathematics skill was also observed. However, it remains unclear the reasons behind hyperconnectivity within these networks. Adopting a Graph Theoretical framework, which analyses network structures and connectivity patterns (please see supplementary information for graph theory metrics’ explanations), could provide valuable insights into the intricate organization of frontoparietal networks and their potential implications for mathematical competency. For example, the excessive edges observed within these specific brain nodes could potentially account for this hyperconnectivity phenomenon (Hillary et al., 2014).

Within the existing literature, the majority of studies investigating the mechanisms of DD or MLD have predominantly employed whole-group analyses, comparing DD or MLD groups to TD groups. However, DD or MLD becomes more intricate upon closer individual scrutiny as they are characterised by heterogeneity on behavioural, cognitive and neural levels (see review from Rubinsten & Henik, 2009). In contrast, there has been a recent surge in attention towards the single-case methodology, introduced by (Crawford & Garthwaite, 2002; Crawford and & Howell, 1998; Crawford et al., 2010), where an individual case of DD or MLD is compared with a group of TD children. This design enables researchers to delve into individual idiosyncrasies, going beyond the limitations imposed by summary statistics derived from group comparisons. Such an approach facilitates the creation of a comprehensive and multi-dimensional taxonomy of DD or MLD (Träff et al., 2017). Furthermore, an in-depth exploration of single cases can provide valuable insights for crafting personalized interventions for children with DD or MLD (Kwasnicka et al., 2019; Wong, 2017).

Given the higher prevalence of MLD compared to DD, and the fact that many low-progress math learners exhibit multiple cognitive deficits, this study aims to deepen the understanding of the underlying neural mechanisms of multiple MLD cases with different deficit profiles in young children. By focusing on MLD rather than exclusively on DD, this approach is more representative of the real-world math learning difficulties encountered in educational settings. Additionally, it addresses the call for more inclusive sampling when investigating math learning difficulties (Astle & Fletcher-Watson, 2020; Astle et al., 2021; Cantlon, 2020).

The current research employed a single-case methodology to investigate brain activations during mathematical tasks and the architectural configurations of frontoparietal networks derived from graph theory during an approximate resting state—when children were instructed to play quietly with toys. The primary objective was to identify underlying neural aberrancies that likely span multiple MLD cases. The key research questions focused on whether aberrant brain activation and network properties could be consistent across these cases. In terms of brain activation, drawing on insights from a similar study (Iuculano et al., 2015), a rough confirmatory hypothesis was drawn that heightened activation in the fronto-parietal areas might be a common feature across all MLD cases. However, due to the scarce of similar research, the possibility of decreased activation was not ruled out. Regarding the frontoparietal network, due to the novel application of graph theory in explaining individual differences among MLD children, it remains challenging to predict which specific aberrant frontoparietal network properties would consistently appear across these cases.

## 2. Materials and Methods

### 2.1 Participants

Sixty-four children in first grade (age: 6.17 – 8.1; M_age_ = 7.08; boys = 37) were recruited across five primary schools. Informed written consent was obtained from the legal guardian of the children and all study protocols were approved by the Institutional Review Board of the Research Integrity and Ethics Office of local university. None of the participants had any other neurological disorders. Only complete and valid data collected from these children were included in the analysis. Ten children were excluded from the study because they met various exclusion criteria outlined in the supplementary document. Data from the remaining fifty-four participants were entered into the final analysis.

### 2.2 Measures

Comprehensive evaluations encompassing both domain-general and domain-specific assessments were conducted by trained researchers, including the Wechsler Individual Achievement Test, Third Edition (WIAT-III; Wechsler, 2009) and the Wechsler Intelligence Scale for Children, Fifth Edition, (WISC-V; Wechsler, 2014). The criteria for identifying individuals within the MLD category were satisfied when children achieved scores below 85 (equivalent to the 16th percentile) on either the Mathematics or Math Fluency subtest of the WIAT-III, and neither of the subtest scores surpassed 90. This criterion is informed by recommendations highlighted in Moeller et al. (2012)’s literature review and Iuculano et al. (2015)’s empirical study. Please see supplementary document—measures—for detailed descriptions of the contents and procedure of these assessments.

### 2.3 Modality and equipment

Neural data was recorded at 8.14 Hz using a continuous wave functional near infra-red spectroscopy system (NIRSport2, NIRx Medical Technologies LLC, Glen Head, NY, United States). FNIRS was one of the most frequently used neuroimaging techniques in educational setting due to its low cost, portable property and robustness to motion, especially in research involving young children (Soltanlou et al., 2018). It infers neural activation from the changes in concentration of oxyhemoglobin (HbO) and deoxygenated hemoglobin (Fekete et al., 2014). In recent years, it was also utilized to investigate the graph theory metrics of brain network (Akila & Johnvictor, 2023; Gallego-Molina et al., 2023; Wang et al., 2019). Thus, it could be used to answer the research question regarding brain activation and graph theory networks.

### 2.4 FNIRS data acquisition

Fifteen LED sources containing 760-nm and 850-nm laser light and fourteen detectors which comprise 39 channels were formed to cover the frontal and parietal areas to HbO changes during resting state, followed by an addition task. The layout of optodes is illustrated in Figure 1 based on the international 10-20 standard system.

**Figure 1.**
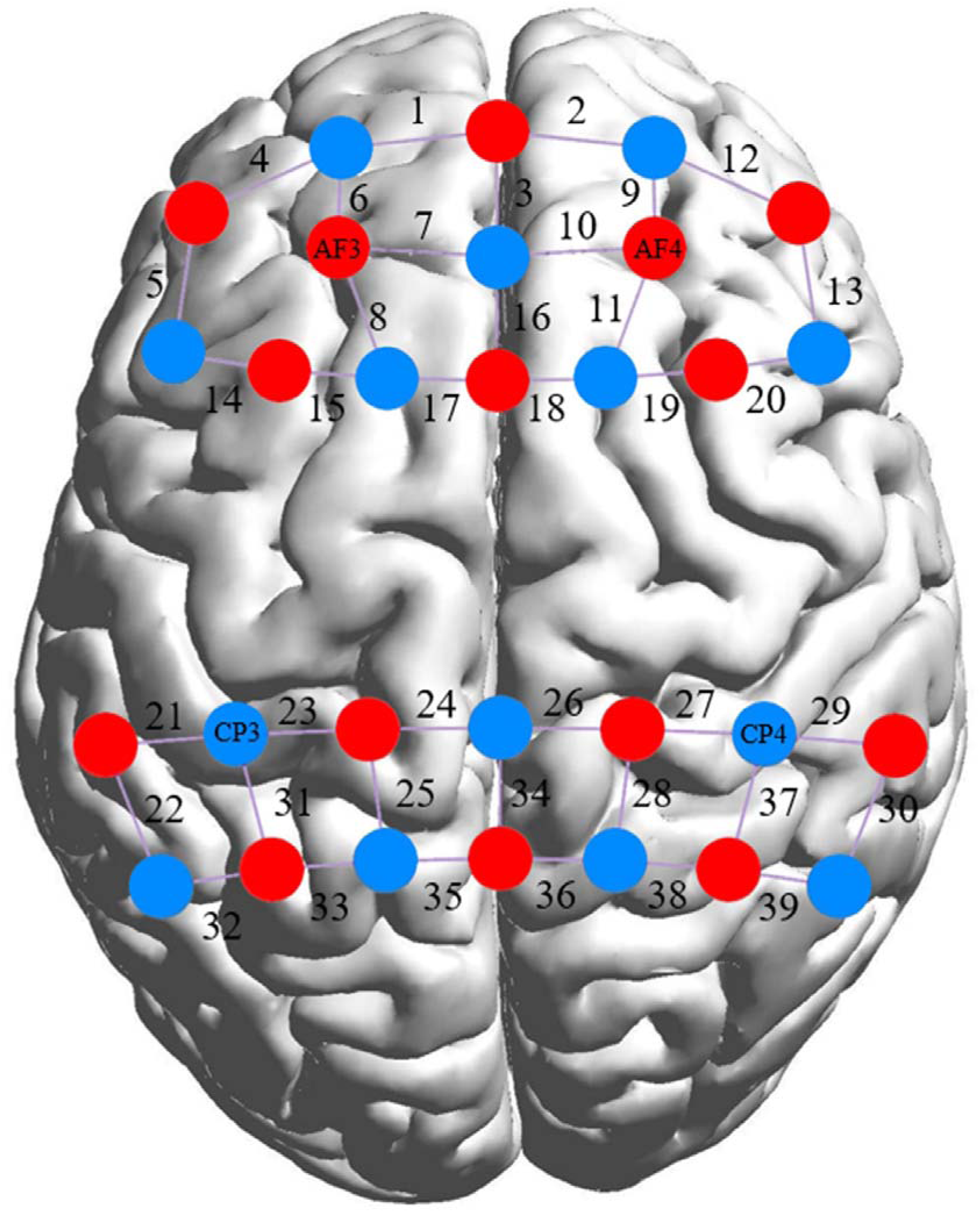
Placement of fNIRS Probes for Freeplay and Addition Task.

### 2.5 Procedure

#### 2.5.1 Procedure for fNIRS approximate resting state

To elicit a condition analogous to the resting state, the research employed the freeplay paradigm introduced by Kim et al. (2020). Within this paradigm, children were afforded the opportunity to engage in unstructured play independently with basic toys (depicted in supplementary document—Figure S4) for a duration of eight minutes.

#### 2.5.2 Procedure for fNIRS experimental task

The experimental protocol was designed to encompass one-digit symbolic and nonsymbolic addition tasks. The task paradigm drew inspiration from three prior neural studies concerning mathematics tasks of varying formats (Gu et al., 2017; Kucian et al., 2011; Venkatraman et al., 2005), with necessary modifications implemented based on the cognitive and developmental demands of the young participants in this study. The sequential progression of the experimental procedure is detailed in Figure 2. It was initiated with a thirty-second fixation phase aimed at establishing a baseline state, during which children were instructed to maintain stillness and direct their gaze towards a cross depicted on the screen. Subsequently, a forty-two-second block dedicated to symbolic or non-symbolic addition tasks, encompassing a series of six trials per block, was presented to the participants. Within each block, children were required to identify the correct answers to a set of addition problems. This task block was succeeded by an eighteen-second rest period. Alternating runs of symbolic and non-symbolic addition tasks were conducted, with a total of four runs per task type, lasting eight minutes in total (Park & Brannon, 2014). The sequence of presenting the symbolic and nonsymbolic blocks was counterbalanced across different participants. Every task block encompassed six individual trials, each lasting for a duration of seven seconds (refer to Figure 2 for an illustration of a symbolic trial and a nonsymbolic trial). Thus, there were forty-eight trails in total. Each trial initiation involved a one-second fixation image featuring a centrally located cross. Following this, a symbolic or nonsymbolic addition problem was presented on the screen for a period of five seconds. During this interval, participants were required to provide responses by pressing the “s” button if the correct answer was situated on the left side of the screen, and the “j” button if the correct answer was positioned on the right side of the screen. This response window was succeeded by a one-second inter-stimulus interval (ISI).

**Figure 2.**
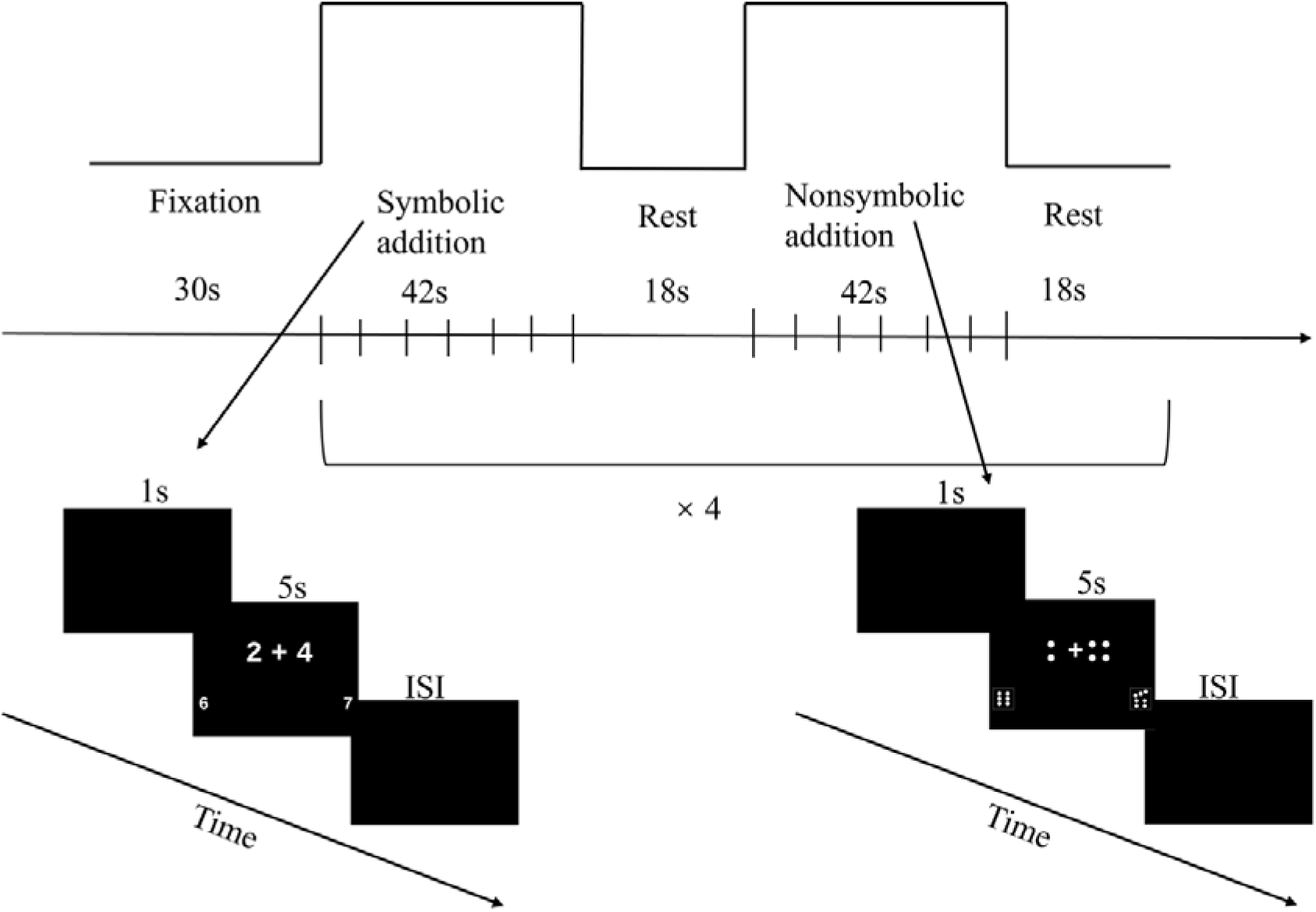
Semantic Flow of Experimental Procedure and A Non/Symbolic Addition Trial.

The operands for symbolic stimuli were Arabic numbers, while nonsymbolic stimuli were presented as dots similar to those on the face of a dice. For all questions, the operands were ranged from 1-5 and problems involving ties (such as 2+2 = 4) were not included (Venkatraman, 2005). For each addition problem, two answer alternatives within the range of 3 to 9 were provided. One alternative was the correct answer, while the other was designed to exhibit a distance effect of one or two units (Pica et al., 2004; Venkatraman, 2005). The assignment of the correct answer to the left or right side of the screen was randomized and balanced across all trial instances. To maintain variety and prevent repetition, the same mathematical problem was never reintroduced within the same task block. The sequence of problems and corresponding answer options were presented in a random fashion for each block. The execution of the experimental paradigm was programmed using E-Prime (E-Prime, Psychology Software Tools Inc. Inc., Pittsburgh, PA).

### 2.6 Data analysis

#### 2.6.1 Task data

The HomER-2 software was used to analyse the fNIRS task-based data (Huppert et al., 2009). First, the signal-to-noise ratio of the data was calculated through the coefficient of variation (CV) within the thirty-nine channels and within each block. Data points were omitted from analysis if the CV for each channel surpassed 15%, and if the CV for each block exceeded 10% for two wavelengths (Van Hedel et al., 2021). The motion artifact was detected by channel in HomER-2. This function found the data-points exceeding a threshold in change of amplitude (1) and a threshold in change of standard deviation (5) within a given period of time (0.5) and then marked those points from the beginning of the window to 1 second later as motion artifacts (Brigadoi et al., 2014). The detected motion artifacts were corrected through Spline interpolation method wherein the degree of Spline function was set to 0.99 as suggested in previous research (Cooper et al., 2012; Scholkmann et al., 2010). Only data between 0.01 and 0.03 Hz were retained for the subsequent steps (Niu et al., 2013). The optical density data was further converted to the concentrations of HbO based on the modified Beer–Lambert law (Delpy et al., 1988; Tak & Ye, 2014). The first five seconds of data within each block were discarded due to unstable fluctuations, resulting in the averaging of thirty-six seconds of data across different blocks.

#### 2.6.2 Approximate resting-state data

The FC-NIRS software (Xu et al., 2015) was used to analyse the approximate resting-state data, with two main procedures to be included: pre-processing and frontoparietal network construction and analysis. The pre-processing procedure comprised quality control, band-pass filters, motion detection and correction, and time selection. First, channels were excluded if the signal-to-noise ratio was less than 2 (Xu et al., 2015). Low-pass filter was set to 0.1 Hz and high-pass filter was set to 0.01 Hz, replicating standards reported in Pelicioni et al. (2019). Motion artifacts were detected through calculating the moving standard deviation within a two-second sliding time window and corrected by Spline interpolation method (Brigadoi et al., 2014). Standard deviation values larger than five standard deviations away from the mean of the moving standard deviation were labelled as artifacts. To construct functional connectivity and frontoparietal network with stability, the analyses excluded the first minute of resting state data acquired and focused on the last seven-minute duration (Wang et al., 2017).

#### 2.6.3 Frontoparietal network construction and analysis

In fNIRS studies, nodes are defined as channels and edges are defined as the functional connectivity between channels (Cai et al., 2019; Qi et al, 2021). Functional connectivity was calculated by performing Pearson correlation analyses between the time series of every pair of nodes for each participant (Wang et al., 2020). Accordingly, a 39 x 39 (channels) symmetric correlation matric was generated for each participant in this study. Subsequently, the Pearson’s R-values were converted to normally distributed z-scores through the function z’ = 0.5*[ln(1 + r) − ln(1–r)] to construct a binary undirected graph (Price et al., 2018; Wang et al, 2018; Zhao et al., 2019). If the correlation value between two nodes equals or exceeds a given threshold, it was set to 1. Otherwise, it was set to 0. The threshold was determined by sparsity, which refers to the ratio of the number of actual edges and the maximum number of edges in a network. The sparsity values ranged from 0.1 to 0.5 with an interval of 0.01 (Xu et al., 2015).

Area under curve was calculated for each network indicator to allow threshold-independent comparisons between different groups (Cai et al., 2019). The normalized clustering coefficient (Gamma) were computed as the ratio of clustering coefficient (Cp) of a real network and mean value of Cp of a randomized network. The normalized characteristic path length (Lambda) was computed as the ratio of characteristic path length (Lp) of a real network and mean value of Lp of a randomized network. 1000 matched random networks that have the same numbers of nodes and edges as the real frontoparietal network were set to calculate the mean normalized clustering coefficient and normalized characteristic path length.

#### 2.6.4 Single-case methodology

Based on case-controls design, the software application “SINGLIMS_ES.exe,” developed (Crawford & Garthwaite, 2002; Crawfordand & Howell, 1998), was employed to conduct a comparative analysis of the distinctions in frontoparietal network configurations and activation patterns among each child exhibiting MLD and the cohort of forty-five TD children. Numerous successful examples of this approach have been documented in the dyscalculia literature, with notable publications including works by Dinkel et al. (2013), Iuculano et al. (2008), Träff et al. (2017), Tressoldi et al. (2007), and Van Viersen et al. (2013), among others. This software enables the comparison of a DD case’s score with criteria derived from a control sample, provided mean and standard deviation of the control sample are available. The algorithm treats the statistics of the normative or control sample as statistical measures rather than population parameters. Additionally, the software employs the t-distribution, instead of the standard normal distribution to gauge the abnormality of individual scores. Essentially, this method resembles a modified independent samples t-test, treating the individual as a sample of N=1 and, consequently, not contributing to the within-group variance estimate (Crawford and & Howell,1998; Crawford & Garthwaite, 2002; Tressoldi et al., 2007). Outcome measures include t-values, p-values and effect size (z_cc_), which indicate an estimate of the average difference between a case’s score and the score of a randomly chosen participant of the normative population (Van Viersen et al., 2013). To address the issue of multiple comparisons, false discovery rate correction (FDR) introduced by Storey (2002), was employed during data analyses of comparing regional nodal characteristics across thirty-nine channels. Only outcomes that remained statistically significant after FDR adjustments were reported.

#### 2.6.5 Group brain- cognition correlational analysis

Although the single-case methodology is the primary analytical approach in this study, group correlational analyses (using Pearson correlation) were also conducted for both groups to provide a more comprehensive understanding of the cognition-brain relationship.

## 3. Results

### 3.1 Demographic information of MLD and TD groups and results of cognitive measures

Out of the study’s recruited participants comprising fifty-four individuals, a subset of nine children satisfied the criteria indicative of being MLD (MLD: 6 boys; mean age = 7.12 years old), while the remaining cohort of forty-five children were categorized as belonging to the TD group (MLD: 21 boys; mean age = 7 years old). This study centred on individual characteristics, evaluating the cognitive performances of nine MLD cases individually. According to the WISC-V criteria, scores between 80-89 fall within the “low average” classification, scores between 70-79 are classified as “well below average,” and scores of 69 and below are categorized as within the “lower extreme” group. Utilizing these benchmarks, a synthesis of the nine cases is presented in Table 1. Notably, despite the diverse constellations of impairments observed among the nine children, a consistent trend emerges in their mathematics-related performance—every case demonstrates performance at least below the average in Mathematics and Math Fluency composites.

**Table 1.** Performance of nine MLD cases with different deficit profiles and classifications.

| Assessment | C1 | C2 | C3 | C4 | C5 | C6 | C7 | C8 | C9 |
| --- | --- | --- | --- | --- | --- | --- | --- | --- | --- |
| Mths | LA<br>(80) | WBA<br>(71) | LA<br>(88) | LE<br>(58) | WBA<br>(78) | LE<br>(65) | WBA<br>(75) | LA<br>(88) | LA<br>(80) |
| MF | LA<br>(81) | WBA<br>(75) | LA<br>(82) | LE<br>(63) | WBA<br>(71) | LE<br>(67) | WBA<br>(76) | LA<br>(81) | A<br>(90) |
| TR | HA<br>(113) | LA<br>(82) | HA<br>(114) | WBA<br>(76) | LE<br>(66) | LE<br>(53) | A<br>(91) | A<br>(94) | LA<br>(87) |
| BR | LA<br>(81) | LE<br>(67) | LA<br>(81) | A<br>(90) | LE<br>(69) | LE<br>(60) | WBA<br>(71) | A<br>(98) | LA<br>(84) |
| WM | WBA<br>(72) | WBA<br>(79) | A<br>(94) | WBA<br>(74) | A<br>(103) | HA<br>(112) | LE<br>(69) | WBA<br>(79) | A<br>(100) |
| FSIQ | WBA<br>(73) | LA<br>(81) | LA<br>(84) | WBA<br>(72) | HA<br>(117) | A<br>(107) | LA<br>(81) | WBA<br>(70) | WBA<br>(73) |

Note: WIAT: Wechsler Individual Achievement Test; Mths: Mathematics; MF: Math Fluency;
TR: Total Reading; BR: Basic Reading; WISC: Wechsler Intelligence Scale for Children;
WM: Working Memory; FSIQ: Full Scale IQ; LA: low average; HA: high average; WBA:
well below average; LE: lower extreme; A: average.

### 3.2 Behavioural results

For behavioural indicators—accuracy and reaction time during (non)symbolic addition tasks, both Case 2 and Case 7 showed significant deviations from the comparison criterion (mean and standard deviation) established by 45 TD children. In particular, Case 2 demonstrated normal accuracy in both symbolic and nonsymbolic tasks but responded significantly faster than TD children in both tasks (see Table 2 for details). On the other hand, Case 7 exhibited impaired abilities across both tasks, with significantly lower accuracy (and marginally significant in nonsymbolic addition) while also responding significantly faster than TD children.

**Table 2.**
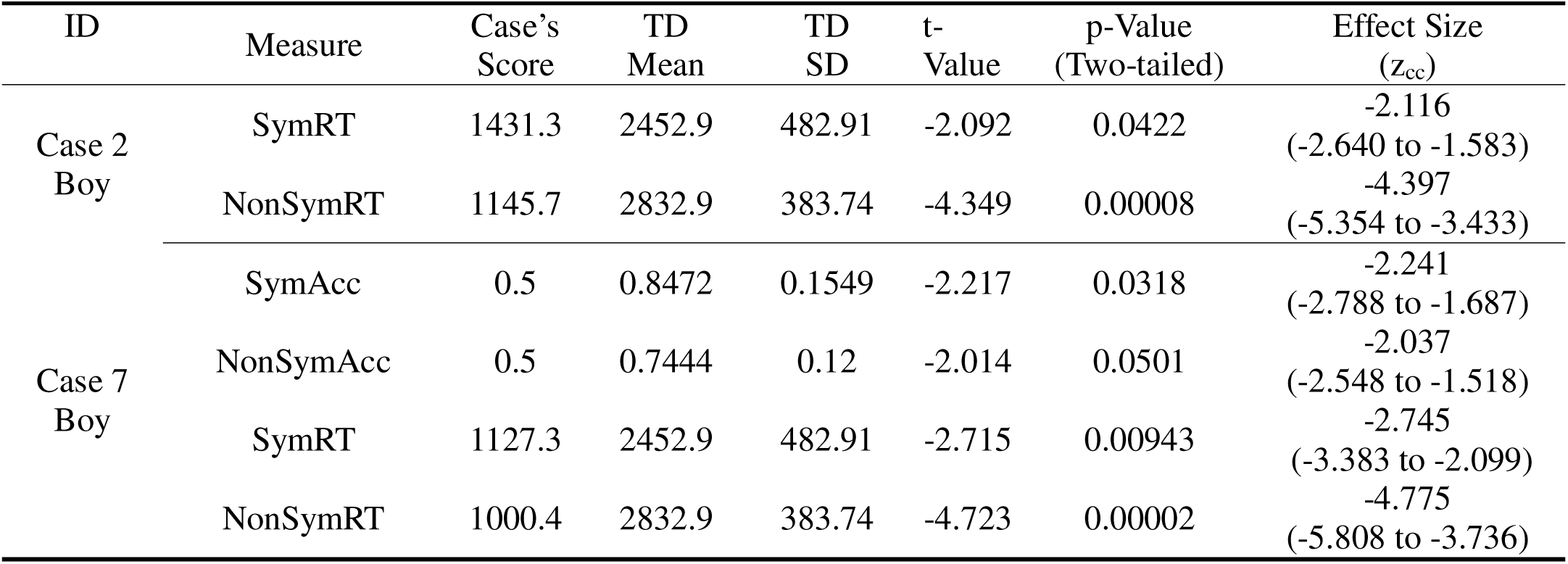
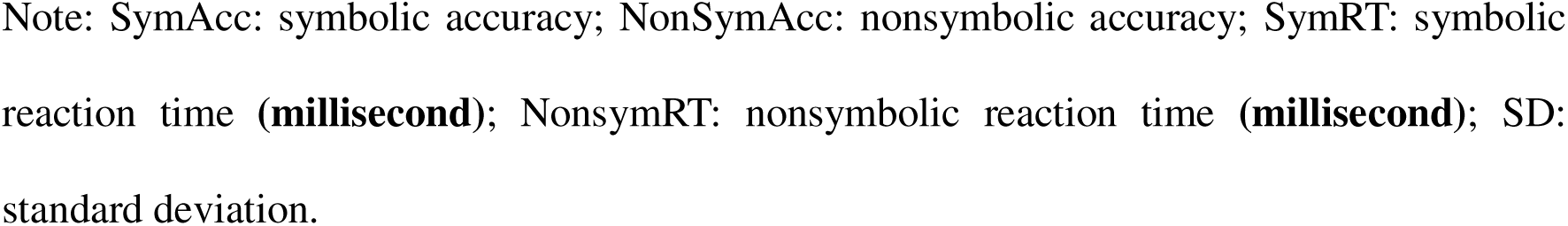
Significantly Different Performance on Neuroimaging Addition Tasks between MLD cases with different deficit profiles and TD Children.

### 3.3 Results of brain activation during addition tasks

Channel-by-channel comparisons were conducted between the nine MLD cases and the TD children on the averaged oxyhaemoglobin concentration in both symbolic and nonsymbolic addition tasks. Only in the symbolic addition task, Case 2 showed significantly decreased activations in the overlapping area of the left inferior and middle frontal gyrus (channel 5) and the left angular gyrus (channel 31). Further particulars are documented in Supplementary Table S2. None of the MLD cases showed aberrant activation in the nonsymbolic addition task.

### 3.4 Results of approximate resting state

Global network properties. Employing single-case methodology, the investigation revealed significant differences between two cases selected from the set of nine MLD cases and the cohort of TD children, particularly in terms of global network properties. One of these cases was identified as Case 2, detailed in supplementary Table S3, while the other case was labelled as Case 3. Case 2 exhibited a significant increase in clustering coefficient, heightened local efficiency, and an extended characteristic path length. On the other hand, Case 3 demonstrated a considerable decrease in global efficiency and a prolonged characteristic path length when compared to the TD children, as documented in Table S3. A salient commonality between these significantly disparate cases is the extended characteristic path length, signifying the minimal number of edges between two nodes.

Nodal network properties. In relation to regional nodal degree properties, it is worth noting that solely two cases, drawn from the potential pool of nine cases, exhibited significantly distinct values in comparison to the forty-five typically developing children. Interestingly, both cases showcased notably reduced nodal degree in comparison to the typically developing children, although in distinct channels. Specifically, Case 3 displayed this phenomenon in right middle frontal gyrus (channel 10), while Case 7 exhibited a decreased nodal degree in the same area but in channel 9, as detailed in supplementary Table S4.

For regional nodal efficiency property, among the three cases aforementioned, a consistent pattern of significantly reduced nodal efficiency in comparison to the TD children was observed, albeit through distinct neural pathways (refer to supplementary Table S5). Case 2 showed a unique profile, with decreased nodal efficiency localized exclusively within channel 27 in the right superior parietal gyrus. Case 3 exhibited decreased nodal efficiency across two extensive neural domains, affecting adjacent channels 1, 3, and 10 in the frontal area, as well as adjacent channels 34, 35, and 36 in the superior parietal area. In contrast, Case 7 displayed a more widespread reduction in nodal efficiency, spanning from the remote left frontal gyrus (channel 4), through the right middle frontal gyrus (channel 9), and extending to the right occipital gyrus (channel 39).

Table 3 outlines the aberrant behavioural and neural indicators exhibited by the three specific cases. Figure 3 shows the locations of the brain areas with inefficient frontoparietal networks in each case. Notably, when employing a single-case methodology, the remaining six MLD cases did not demonstrate any aberrations in comparison to TD children.

**Figure 3.**
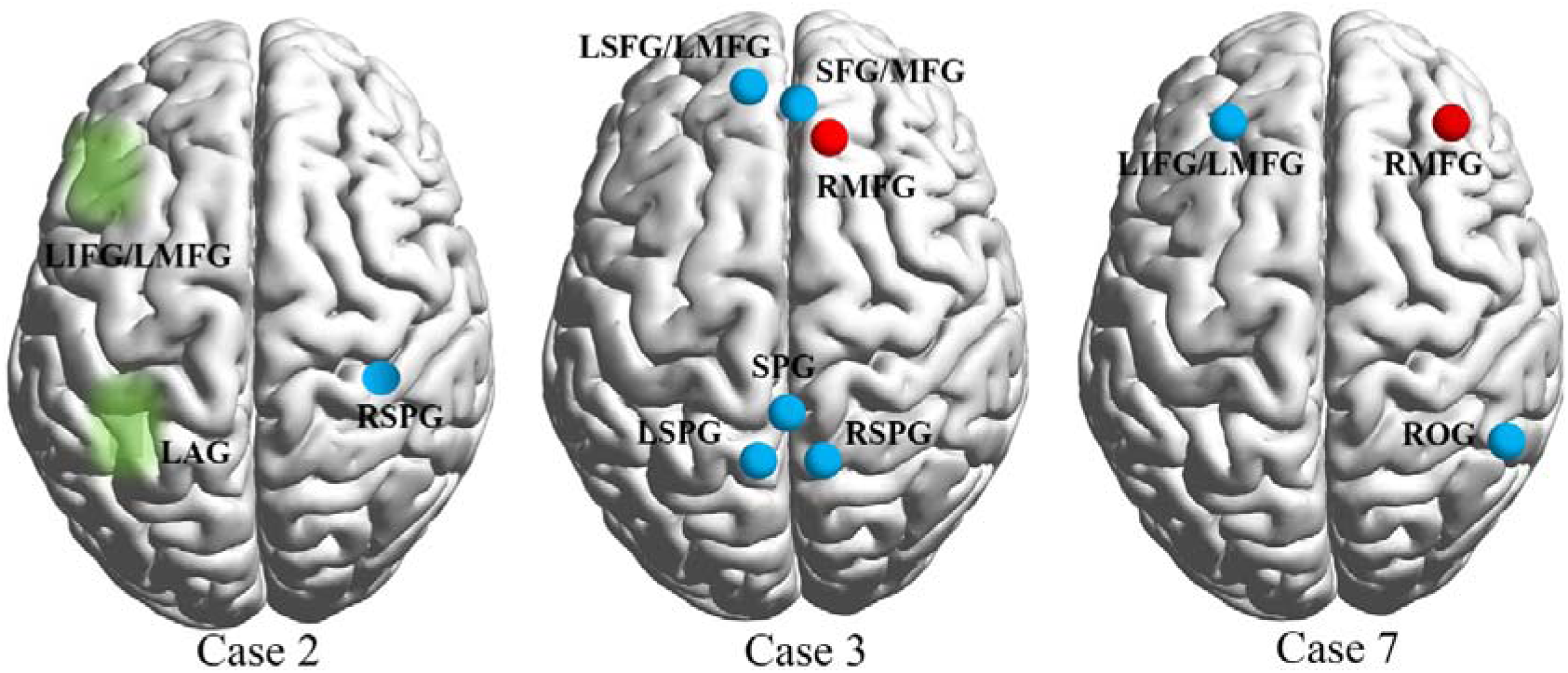
The Abnormal Regional Network Properties and Activation of Three Cases Showed.

**Table 3.** Summary of the Aberrant Behavioural and Neural Indicators.

| Case ID | Task-based behavioural indicators | Task-based Brain Activation | Resting-state Global Properties | Resting-state Nodal Properties |
| --- | --- | --- | --- | --- |
| 2, boy | Symbolic and nonsymbolic reaction time | Lower activation in symbolic addition (ch5, ch31) | Longer characteristic path length, higher clustering coefficient and higher local efficiency | Lower nodal efficiency (ch27) |
| 3, girl | Not Applicable | Not Applicable | Lower global efficiency; longer characteristic path length | Lower nodal degree (ch10); Lower nodal efficiency (ch1, 3, 10, 34, 35, 36). |
| 7, boy | (non)symbolic accuracy and reaction time | Not Applicable | Not Applicable | Lower nodal degree (ch9), Lower nodal efficiency (ch 4, 9, 39) |

**Note:** ch1: Left superior/middle frontal gyrus; ch 3: Superior/middle frontal gyrus; ch4: Left middle/inferior frontal gyrus; ch 5: Left middle/inferior frontal gyrus; ch 9: Right middle frontal gyrus; ch10: Right middle frontal gyrus; ch27: Right superior parietal gyrus; ch31: Left angular gyrus; ch34: superior parietal gyrus; ch35: Left superior parietal gyrus; ch36: Right superior parietal gyrus; ch39: Right occipital gyrus.

### 3.5 Group brain-cognition correlational results

Only the nodal efficiency on channel 4, the junction between the left inferior and middle frontal gyrus for TD children was positively related to math fluency (*r* = 0.31, *p* = 0.046) and accuracy in nonsymbolic addition task (*r* = 0.329, *p* = 0.033) while there was no significant correlation between other cognitive performances and neural indicators. The positive correlation results indicated that math performance would slightly be improved with higher nodal efficiency within this junction as Figure 4 shows.

**Figure 4.**
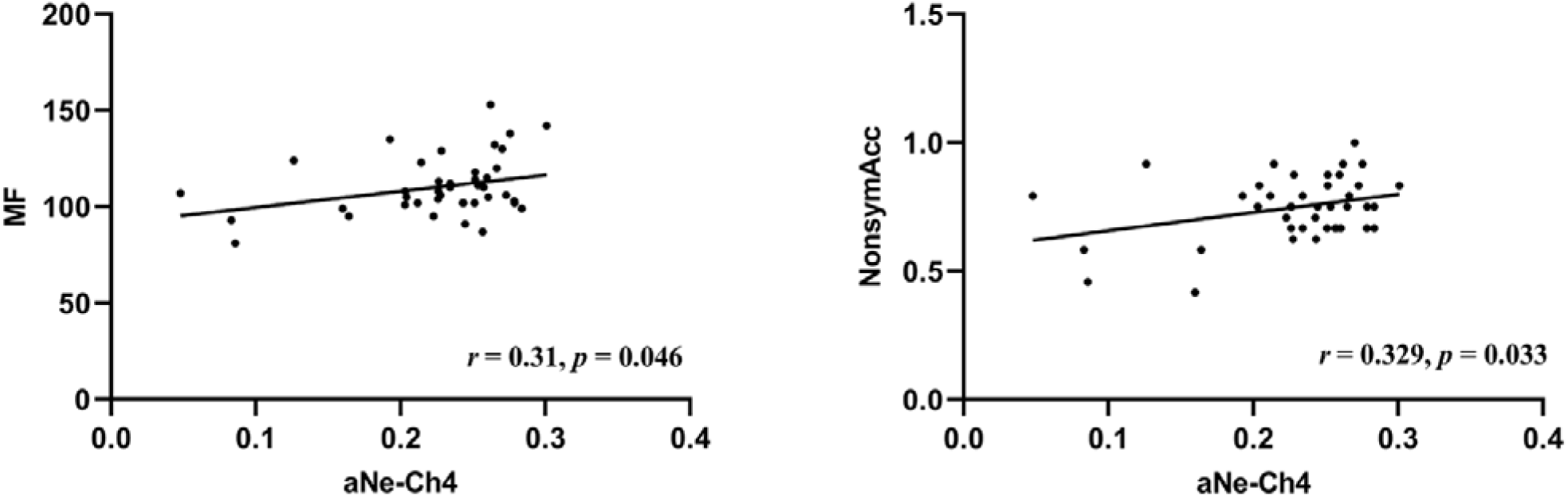
Correlations between Nodal Efficiency of Channel 4 and Math fluency and Accuracy of Nonsymbolic Addition in TD Children.

## 4. Discussion

This study sought to identify underlying neural aberrancies that likely span multiple MLD cases with heterogenous cognitive profiles. The comprehensive findings revealed that there were no unified aberrancies spanning all MLD cases, regardless of brain activation or network properties. The discussion will begin with a focus on the three cases that displayed aberrant neural indicators, compared to TD children. Following this, emphasis will be placed on the remaining six cases where no neural.

### 4.1 Brain activation differences between MLD cases and TD children

It was hypothesized that increased brain activation in the frontal-parietal areas would be a common feature across all cases of MLD. However, this pattern was not observed in any of the cases. Specifically, Case 2 exhibited significantly lower brain activation in channels 5 and 31 compared to TD children. Channel 5 is located at the junction of the left inferior and middle frontal gyrus, while Channel 31 is situated in the left angular gyrus.

A similar study by Iuculano et al. (2015) using fMRI during single-digit addition tasks found that TD children exhibited significantly higher brain activation in the superior frontal gyri and the inferior/superior parietal cortices, including the left IPS. As highlighted in the Introduction, a statistically significant difference between groups does not necessarily imply that all individuals in the experimental group exhibit traits entirely distinct from those in the control group, and vice versa (Tressoldi et al., 2007). In this single-case study, only Case 2 demonstrated decreased activation in both the left inferior/middle frontal gyrus and the left angular gyrus.

Although this study employs a similar task and the same cut-off inclusion percentage for MLD as Iuculano et al. (2015), there is a notable difference in participant age. In the current study, participants were 6-8 years old and in the first grade, whereas participants in Iuculano et al.’s study were 7-9 years old and in the third grade. Literature suggests that a shift in activation during addition tasks from prefrontal areas in children to parietal areas in adults, indicating a reduced reliance on working memory with maturation (Rivera et al., 2005). However, it is unclear whether a similar shift from inferior to superior activation in the frontal areas occurs during the early school years, leaving this as an open question for future research.

In Iuculano et al.’s study, the higher activation in the MLD group was interpreted as over-engagement. In contrast, the lower activation observed in Case 2 of the current study might suggest under-engagement. This finding could indicate that Case 2 struggled to effectively engage the left inferior/middle frontal gyrus and the left angular gyrus, which are critical for successful arithmetic processing (Berteletti et al., 2014). One possible explanation is that Case 2’s difficulties in utilizing these essential areas for efficient mental calculations led to a notably quicker completion of the symbolic addition task, as shown in Table 2. This expedited completion may reflect his challenges in engaging the brain regions typically involved in the more time-consuming process of mental arithmetic, which relies on a range of cognitive functions.

Specifically, the left inferior/middle frontal gyrus is associated with semantic processing (Devlin et al., 2003; Poldrack et al., 1999), language processing (Gabrieli et al., 1998; Zhang et al., 2004), and working memory (Baldo & Dronkers, 2006; Öztekin et al., 2009). Upon reviewing the cognitive profile of Case 2, the individual exhibited characteristics of dyslexia as well as working memory deficits. The reduced activation in this brain region corresponds with previous findings of atypical neural activity observed in children with dyslexia during language processing and reading tasks (Kwok & Ansari, 2019). The left angular gyrus, on the other hand, is linked to fact retrieval in mental arithmetic (De Smedt, 2016; Smaczny et al., 2021; Sokolowski et al., 2023). While under-engagement could also be interpreted as a sign of a more automatic or skilled performance, corroborating this with Case 2’s low arithmetic accuracy makes it unlikely that the lower brain activation is due to an automatic brain response associated with superior performance.

### 4.2 Frontoparietal network differences between MLD cases and TD children

Regarding the frontoparietal network, there were no aberrancies spanning all MLD cases. Only Case 2, 3 and 7 showed aberrant frontoparietal network, compared to TD children. The only similarity across these three cases was the lower nodal efficiency, but in different brain areas.

Specifically, Case 2 exhibited both a higher clustering coefficient and local efficiency, where a higher clustering coefficient indicates that more neighbouring nodes are connected to each other, thus correlating with a higher local efficiency. Both a higher clustering coefficient and higher local efficiency indicated that his frontoparietal network orient towards a cliquish (i.e., a pattern in which certain groups of brain regions tend to be more interconnected with each other than with regions outside of those groups) organization of the brain (Klados, et al., 2013). While no studies have applied graph theory to investigate MLD, one study by Gallego-Molina et al. (2023) combined fNIRS data and graph theory to examine children with dyslexia. They found that dyslexic children exhibited a larger clustering coefficient compared to TD children, which was interpreted as reflecting difficulties in quickly settling into phonological representations. Similarly, the larger clustering coefficient observed in Case 2 who also showed signs of dyslexia may indicate a highly interconnected frontoparietal network, potentially limiting his flexibility in allocating brain resources during mathematical tasks (Jolles et al., 2016; Iuculano, 2016).

Additionally, Case 2 in channel 27 and Case 3 in channel 34, 35 and 36 displayed lower nodal efficiency, which are all located in the superior parietal gyri. These areas are considered relevant to mathematics in six-year-old children (Schel & Klingberg, 2017). To date, the existing body of literature has primarily centred around the activation patterns of the bilateral superior parietal gyri during tasks related to mathematics, while simultaneously engaging in debates regarding the strength of their activations. This present study contributes additional insights derived from the properties of the superior parietal gyrus as revealed by Graph Theory analysis. Successful mathematics performance relies on their interaction with other brain areas (Iuculano & Menon, 2018; Jolles et al., 2016; Menon & Chang, 2021). Should their efficiency in processing information with other distant brain regions be compromised, it could lead to an inability to effectively orchestrate interactions between the inferior frontal gyrus and fusiform gyrus during mathematical tasks. This finding complemented the role of superior parietal gyrus in its pivotal topological organization, which may facilitate mathematical tasks.

In the case of Case 3, she exhibited a longer characteristic path length and lower global efficiency in global network properties. It is important to note that characteristic path length and global efficiency are inversely related by their formulas. The reduction in global efficiency observed in her case was consistent with findings from research on varying complexity levels of mathematical tasks (Klados et al., 2013). This research showed lower global efficiency during less complex tasks, suggesting that individuals with reduced global efficiency may demonstrate less optimal brain functioning.

Specifically, Case 7 exhibited diminished nodal degree in right middle frontal gyrus (channel 9), coupled with reduced nodal efficiency in the junction of left inferior and middle frontal gyrus (channel 4), right middle frontal gyrus (channel 9), and right occipital gyrus (channel 39). Despite these local discrepancies, his global network properties did not exhibit significant differences when compared to those of TD children. In Case 7, lower nodal efficiency in channel 4, located at the junction of the left and middle inferior gyrus, aligns with the group correlational analysis of TD individuals between the nodal efficiency of this junction and mathematics performance. Specifically, the TD group showed a positive correlation between nodal efficiency at this junction and both Math Fluency and Accuracy in Nonsymbolic Addition. These findings suggest that higher nodal efficiency at this junction is associated with better mathematics performance. Higher nodal efficiency indicates that a node, on average, connects to all other nodes in the network via shorter paths, which generally facilitates faster signal transmission with lower energy costs (Bullmore & Sporns, 2012).

Given the limited research on graph theory metrics in children with MLD, this finding was discussed in relation to a study on intelligence and nodal efficiency (Hilger et al., 2017). Mathematical performance is generally closely linked to intelligence (Song & Su, 2022). It is important to note that, in addition to MLD, Case 7 also demonstrated a low-average IQ. Hilger et al. found that higher IQ was positively related to greater nodal efficiency in the brain’s salience network. Although the salience network does not include this specific junction, these results suggest a broader relationship between nodal efficiency and cognitive performance, including mathematics. In addition to its role in activation during math tasks, as discussed in Case 2, the current findings support the idea that this junction contributes to individual differences in brain efficiency.

Notably, in Case 3, channel 10, and Case 7, channel 9, both correspond to the right middle frontal gyrus. This region exhibited both decreased degree and efficiency in these cases, potentially indicating weakened connections and reduced efficiency in information transfer with other brain regions. It is plausible that this area functioned as a “hub region” in this Case, a node or group of nodes within a network characterized by high connectivity (Van den Heuvel & Sporns, 2013). When a hub region is compromised, it can lead to more significant alterations in the overall network architecture and information transfer efficiency compared to non-hub regions (Aerts et al., 2016).

### 4.3 No differences supported by the other six MLD cases

Despite the three cases displaying significantly aberrant behavioural and/or neural indicators, these indicators in other six MLD cases were within the normal range of TD cases. The existing literature has delineated a range of compensatory mechanisms aimed at elucidating the factors contributing to neural performance parity with typically developing children. These mechanisms encompass the recruitment of additional brain regions supportive of domain-general abilities, coupled with potential heightened activations within regions specialized for numerical processing in contrast to typically developing peers (Fengjuan & Jamaluin, 2023). It is essential to acknowledge, however, that the placement of fNIRS probes in the present study only covers a portion of the frontal and parietal regions. Consequently, there exists a possibility that compensatory mechanisms may occur in those regions left uncovered by the probes.

It is also important to note that the single-case methodology used in this study tends to produce a more conservative t-statistic compared to conventional approaches like the z-test or independent-samples t-test (Ferreira-Correia & Cockcroft, 2023; Iuculano et al., 2008). While the performance of the six cases appears to fall within the normal range based on this methodology, this does not necessarily indicate normality in their behavioral and neural performances when assessed using other, potentially more lenient, methods. Furthermore, this approach does not rule out the possibility that these cases may possess other “aberrant brain properties” that remain undetected by the current methodologies. As Astle and Fletcher-Watson highlighted in their review (2020), many children who struggle with learning difficulties may never receive a formal diagnosis, even though they meet the criteria for multiple different conditions.

On the other hand, the findings from the six cases align with two recent studies, which suggest that groups with MLD or DD do not differ from TD groups at either the neural or behavioural level. Kwok et al. (2023) conducted univariate and frequentist fMRI analyses on 30 children with DD and 38 TD children, finding no group differences in brain activation during tasks involving arithmetic, magnitude processing, and visuospatial working memory. Although behaviourally, children with DD were less accurate than their typically developing peers in basic number processing and arithmetic tasks, this did not correspond to atypical brain activation at the neural level. From a behavioural perspective, Mammarella et al. (2021) tested both real MLD children and a simulated population to investigate the concept of a core deficit in MLD. They found that none of the measures of basic number processing or domain-general abilities could identify a ‘core deficit’ in children with MLD. This new evidence, together with the six cases in this study, contradicts previous literature suggesting that children with MLD differ from TD children in behavioural and/or neural indicators. With limited evidence supporting this emerging perspective, it remains challenging to overturn the long-standing conclusions about “group differences”. However, advanced methodologies, including multivariate, network-based approaches, and simulation techniques, may allow future research to further explore and test this novel conclusion.

This study did not identify a unified brain aberrancy that spans across all MLD participants. Different cognitive profiles, such as above-average working memory, reading ability, and IQ, as shown in Table 1, may partially mitigate MLD. This underscores the idea that MLD should not be seen as a purely isolated learning “disorder,” but rather as a condition shaped by the complex interactions among various cognitive abilities. Additionally, the inconsistency in findings across individual cases highlights the limitations of relying on group comparison. Even if a group of children with MLD is found to differ from TD children, it does not guarantee that each MLD case will show aberrations in the measures of interest.

The identification of characteristics that consistently span across all children with MLD remains one of the most recent and advanced hypotheses in this field (Astle et al., 2021; Astle & Fletcher-Watson, 2020). In this study, the only potential indicator identified was lower nodal efficiency, which may be linked to poorer mathematical performance. We anticipate that future research will further validate this finding, suggesting that, while not all children with MLD may have impaired nodal efficiency, a ratio of around 33% (3 out of 9 in this study) is likely. It is advisable to combine the complementary strengths of both group comparison with single-case methodology to uncover the underlying neural mechanisms common to all children with MLD.

## Limitations

This study has two main limitations. First, while free play was used to approximate a resting state due to the participants’ young age, it does not fully capture a true resting state in which no cognitive tasks are engaged. This should be taken into account when interpreting the findings. Second, although the case-control design provided a detailed understanding of the unique characteristics of MLD cases with different deficit profiles, the results should not be generalized to explain the neural mechanisms of MLD in its entirety. The heterogeneity among MLD cases, reflected in the three cases with distinct neural aberrancies, suggests that each case may represent a specific subgroup within the broader MLD population. Future large-scale studies are needed to explore these subgroups further and clarify their underlying neural mechanisms.

## Acknowledgements

This research was funded by the Singapore National Research Foundation (NRF) under the Science of Learning Initiative (NRF2016-SOL002-003) and Nanyang Technological University (NTU) project (SUG-NAP 8/18 JA). Wang Fengjuan is supported by NTU Research Scholarship. Any opinions, findings, and conclusions or recommendations expressed in this material are those of the author(s) and do not necessarily reflect the views of the NRF or NIE/NTU.

## Abbreviations

MLD: mathematics learning difficulties
DD: developmental dyscalculia
TD: typically developing
fNIRS: functional near-infrared spectroscopy
RSFC: resting-state functional connectivity
IPS: intraparietal sulcus
HbO: oxyhemoglobin
ISI: inter-stimulus interval
CV: coefficient of variation
FDR: false discovery rate correction
WIAT-III: Wechsler Individual Achievement Test, Third Edition
WISC-V: Wechsler Intelligence Scale for Children, Fifth Edition
Mths: Mathemati
MF: Math Fluency
TR: Total Reading
BR: Basic Reading
WISC: Wechsler Intelligence Scale for Children
WM: Working Memory
FSIQ: Full Scale IQ
LA: low average
HA: high average
WBA: well below average
LE: lower extreme
A: average
SD: standard deviation
aNe-Ch4: nodal efficiency of channel 4

## Data availability statement

The data that support the findings of this study will be openly available in Figshare.

## References

Aerts, H., Fias, W., Caeyenberghs, K., & Marinazzo, D. (2016). Brain networks under attack: robustness properties and the impact of lesions. Brain, 139(12), 3063–3083. 10.1093/brain/aww194

Akila, V., & Johnvictor, A. C. (2023). Functional near infrared spectroscopy for brain functional connectivity analysis: A graph theoretic approach. Heliyon, 9(4). 10.1016/j.heliyon.2023.e15002

American Psychiatric Association (2013). Diagnostic and Statistical Manual of Mental Disorders. Arlington, VA: American Psychiatric Association.

Astle, D. E., & Fletcher-Watson, S. (2020). Beyond the core-deficit hypothesis in developmental disorders. Current Directions in Psychological Science, 29(5), 431–437. 10.1177/0963721420925518

Astle, D. E., Holmes, J., Kievit, R., & Gathercole, S. E. (2022). Annual Research Review: The transdiagnostic revolution in neurodevelopmental disorders. Journal of Child Psychology and Psychiatry, 63(4), 397–417. 10.1111/jcpp.13481

Baldo, J. V., & Dronkers, N. F. (2006). The role of inferior parietal and inferior frontal cortex in working memory. Neuropsychology, 20(5), 529. https://psycnet.apa.org/doi/10.1037/0894-4105.20.5.529

Berteletti, I., Prado, J., & Booth, J. R. (2014). Children with mathematical learning disability fail in recruiting verbal and numerical brain regions when solving simple multiplication problems. Cortex, 57, 143–155. 10.1016/j.cortex.2014.04.001

Biswal, B., Zerrin Yetkin, F., Haughton, V. M., & Hyde, J. S. (1995). Functional connectivity in the motor cortex of resting human brain using echo-planar MRI. Magnetic resonance in medicine, 34(4), 537–541. 10.1002/mrm.1910340409

Bullmore, E., & Sporns, O. (2012). The economy of brain network organization. Nature Reviews. Neuroscience, 13(5), 336–349. 10.1038/nrn3214

Butterworth, B., Varma, S., & Laurillard, D. (2011). Dyscalculia: from brain to education. science, 332(6033), 1049–1053. 10.1126/science.1201536

Butterworth, B. (2019). Dyscalculia: From science to education. Routledge.

Brigadoi, S., Ceccherini, L., Cutini, S., Scarpa, F., Scatturin, P., Selb, J., & Cooper, R. J. (2014). Motion artifacts in functional near-infrared spectroscopy: a comparison of motion correction techniques applied to real cognitive data. Neuroimage, 85, 181–191. 10.1016/j.neuroimage.2013.04.082

Cai, L., Dong, Q., Wang, M., & Niu, H. (2019). Functional near-infrared spectroscopy evidence for the development of topological asymmetry between hemispheric brain networks from childhood to adulthood. Neurophotonics, 6(2), 025005–025005. 10.1117/1.NPh.6.2.025005

Cantlon, J. F. (2020). The balance of rigor and reality in developmental neuroscience. NeuroImage, 216, 116464. 10.1016/j.neuroimage.2019.116464

Cooper, R. J., Selb, J., Gagnon, L., Phillip, D., Schytz, H. W., Iversen, H. K., … & Boas, D. A. (2012). A systematic comparison of motion artifact correction techniques for functional near-infrared spectroscopy. Frontiers in neuroscience, 6, 147. 10.3389/fnins.2012.00147

Crawford, J. R., & Garthwaite, P. H. (2002). Investigation of the single case in neuropsychology: Confidence limits on the abnormality of test scores and test score differences. Neuropsychologia, 40, 1196–1208. 10.1016/S0028-3932(01)00224-X

Crawford, J. R., Garthwaite, P. H., & Wood, L. T. (2010). Inferential methods for comparing two single cases. Cognitive neuropsychology, 27(5), 377–400. 10.1080/02643294.2011.559158

Crawford, J. R., & Howell, D. C. (1998). Comparing an individual’s test score against norms derived from small samples. The Clinical Neuropsychologist, 12, 482–486. 10.1076/clin.12.4.482.7241

De Smedt, B. (2016). Individual differences in arithmetic fact retrieval. In Development of mathematical cognition (pp. 219–243). Academic Press. 10.1016/B978-0-12-801871-2.00009-5

Delpy, D. T., Cope, M., van der Zee, P., Arridge, S., Wray, S., & Wyatt, J. S. (1988). Estimation of optical pathlength through tissue from direct time of flight measurement. Physics in Medicine & Biology, 33(12), 1433. DOI 10.1088/0031-9155/33/12/008

Devlin, J. T., Matthews, P. M., & Rushworth, M. F. (2003). Semantic processing in the left inferior prefrontal cortex: a combined functional magnetic resonance imaging and transcranial magnetic stimulation study. Journal of cognitive neuroscience, 15(1), 71–84. 10.1162/089892903321107837

Dinkel, P. J., Willmes, K., Krinzinger, H., Konrad, K., & Koten Jr, J. W. (2013). Diagnosing developmental dyscalculia on the basis of reliable single case FMRI methods: promises and limitations. PLoS One, 8(12), e83722. 10.1371/journal.pone.0083722

Fekete, T., Beacher, F. D. C. C., Cha, J., Rubin, D., & Mujica-Parodi, L. R. (2014). Small-world network properties in prefrontal cortex correlate with predictors of psychopathology risk in young children: A NIRS study. NeuroImage, 85, 345–353. 10.1016/j.neuroimage.2013.07.022

Fengjuan, W., Jamaludin, A. (2023). The Science of Mathematics Learning: An Integrative Review of Neuroimaging Data in Developmental Dyscalculia. In: Hung, W.L.D., Jamaludin, A., Rahman, A.A. (eds) Applying the Science of Learning to Education. Springer, Singapore. 10.1007/978-981-99-5378-3_3

Ferreira-Correia, A., & Cockcroft, K. (2023). Controlling for inequality in neuropsychological assessment: using Crawford and Howell’s (1998) single-case methodology with norms from demographically homogeneous groups of South Africans. South African Journal of Psychology, 53(3), 327–340. 10.1177/00812463221151008

Gabrieli, J. D., Poldrack, R. A., & Desmond, J. E. (1998). The role of left prefrontal cortex in language and memory. Proceedings of the national Academy of Sciences, 95(3), 906–913. 10.1073/pnas.95.3.906

Gallego-Molina, N. J., Ortiz, A., Martínez-Murcia, F. J., Rodríguez-Rodríguez, I., & Luque, J. L. (2023). Assessing functional brain network dynamics in dyslexia from FNIRS data. International Journal of Neural Systems, 33(04), 2350017. 10.1142/S012906572350017X

Gilmore, C., Clayton, S., Cragg, L., McKeaveney, C., Simms, V., & Johnson, S. (2018). Understanding arithmetic concepts: The role of domain-specific and domain-general skills. PloS one, 13(9), e0201724. 10.1371/journal.pone.0201724

Gu, Y., Miao, S., Han, J., Zeng, K., Ouyang, G., Yang, J., & Li, X. (2017). Complexity analysis of fNIRS signals in ADHD children during working memory task. Scientific Reports, 7(1), 1–10. 10.1038/s41598-017-00965-4

Hawes, Z., Sokolowski, H. M., Ononye, C. B., & Ansari, D. (2019). Neural underpinnings of numerical and spatial cognition: An fMRI meta-analysis of brain regions associated with symbolic number, arithmetic, and mental rotation. Neuroscience & Biobehavioral Reviews, 103, 316–336. 10.1016/j.neubiorev.2019.05.007

Hilger, K., Ekman, M., Fiebach, C. J., & Basten, U. (2017). Efficient hubs in the intelligent brain: Nodal efficiency of hub regions in the salience network is associated with general intelligence. Intelligence, 60, 10–25. 10.1016/j.intell.2016.11.001

Hillary, F. G., Rajtmajer, S. M., Roman, C. A., Medaglia, J. D., Slocomb-Dluzen, J. E., Calhoun, V. D., … & Wylie, G. R. (2014). The rich get richer: brain injury elicits hyperconnectivity in core subnetworks. PloS one, 9(8), e104021. 10.1371/journal.pone.0104021

Huppert, T. J., Diamond, S. G., Franceschini, M. A., & Boas, D. A. (2009). HomER: a review of time-series analysis methods for near-infrared spectroscopy of the brain. Applied Optics, 48(10), D280–D298. 10.1364/AO.48.00D280

Iuculano, T. (2016). Neurocognitive accounts of developmental dyscalculia and its remediation. Progress in brain research, 227, 305–333. 10.1016/bs.pbr.2016.04.024

Iuculano, T., & Menon, V. (2018). Development of mathematical reasoning. *Stevens’* Handbook of Experimental Psychology and Cognitive Neuroscience, 4, 183–222.

Iuculano, T., Rosenberg-Lee, M., Richardson, J., Tenison, C., Fuchs, L., Supekar, K., & Menon, V. (2015). Cognitive tutoring induces widespread neuroplasticity and remediates brain function in children with mathematical learning disabilities. Nature Communications, 6(1), 1–10. 10.1038/ncomms9453

Iuculano, T., Tang, J., Hall, C. W., & Butterworth, B. (2008). Core information processing deficits in developmental dyscalculia and low numeracy. Developmental science, 11(5), 669–680. 10.1111/j.1467-7687.2008.00716.x

Jolles, D., Ashkenazi, S., Kochalka, J., Evans, T., Richardson, J., Rosenberg-Lee, M., Zhao, H., Supekar, K., Chen, T., & Menon, V. (2016). Parietal hyperconnectivity, aberrant brain organization, and circuit-based biomarkers in children with mathematical disabilities. Developmental Science, 19(4), 613–631. 10.1111/desc.12399

Kim, J., Ruesch, A., Kang, N. R., Huppert, T. J., Kainerstorfer, J., Thiessen, E. D., & Fisher, A. V. (2020). A Paradigm for Measuring Resting State Functional Connectivity in Young Children Using fNIRS and Freeplay. bioRxiv, 2020-01. doi: 10.1101/2020.01.13.904029

Klados, M. A., Kanatsouli, K., Antoniou, I., Babiloni, F., Tsirka, V., Bamidis, P. D., & Micheloyannis, S. (2013). A graph theoretical approach to study the organization of the cortical networks during different mathematical tasks. PLoS One, 8(8), e71800. 10.1371/journal.pone.0071800

Kwok, F. Y., & Ansari, D. (2019). The promises of educational neuroscience: examples from literacy and numeracy. Learning: Research and Practice, 5(2), 189–200.

Kucian, K., Loenneker, T., Martin, E., & von Aster, M. (2011). Non-symbolic numerical distance effect in children with and without developmental dyscalculia: A parametric fMRI study. Developmental Neuropsychology, 36(6), 741–762. 10.1080/87565641.2010.549867

Kwasnicka, D., Inauen, J., Nieuwenboom, W., Nurmi, J., Schneider, A., Short, C. E., … & Naughton, F. (2019). Challenges and solutions for N-of-1 design studies in health psychology. Health Psychology Review, 13(2), 163–178. 10.1080/17437199.2018.1564627

Kwok, F. Y., Wilkey, E. D., Peters, L., Khiu, E., Bull, R., Lee, K., & Ansari, D. (2023). Developmental dyscalculia is not associated with atypical brain activation: A univariate fMRI study of arithmetic, magnitude processing, and visuospatial working memory. Human Brain Mapping, 44(18), 6308–6325. 10.1002/hbm.26495

Luoni, C., Scorza, M., Stefanelli, S., Fagiolini, B., & Termine, C. (2023). A neuropsychological profile of developmental dyscalculia: the role of comorbidity. Journal of Learning Disabilities, 56(4), 310–323. 10.1177/00222194221102925

Mammarella, I. C., Toffalini, E., Caviola, S., Colling, L., & Szűcs, D. (2021). No evidence for a core deficit in developmental dyscalculia or mathematical learning disabilities. Journal of Child Psychology and Psychiatry, 62(6), 704–714. 10.1111/jcpp.13397

Mazzocco, M. M. M. (2007). Defining and differentiating mathematical learning disabilities and difficulties. In D. B. Berch & M. M. M. Mazzocco (Eds.), Why is math so hard for some children? The nature and origins of mathematical learning difficulties and disabilities (pp. 29–47). Paul H. Brookes Publishing, Baltimore, MD, US

Menon, V. (2016). Working memory in children’s math learning and its disruption in dyscalculia. Current Opinion in Behavioral Sciences, 10, 125–132. 10.1016/j.cobeha.2016.05.014

Menon, V., & Chang, H. (2021). Emerging neurodevelopmental perspectives on mathematical learning. Developmental Review, 60, 100964. 10.1016/j.dr.2021.100964

Michels, L., O’Gorman, R., & Kucian, K. (2018). Functional hyperconnectivity vanishes in children with developmental dyscalculia after numerical intervention. Developmental Cognitive Neuroscience, 30, 291–303. 10.1016/j.dcn.2017.03.005

Ministry of Education. 2020, November, 2. Percentage of PLSE Students Who Scored A*-C, By Subject. https://data.gov.sg/dataset/percentage-of-psle-students-who-scored-a-c-by-subject

Moeller, K., Fischer, U., Cress, U., & Nuerk, H. C. (2012). Diagnostics and intervention in developmental dyscalculia: Current issues and novel perspectives (pp. 233–275). Springer Netherlands.

Niu, H., Li, Z., Liao, X., Wang, J., Zhao, T., Shu, N., Zhao, X., & He, Y. (2013). Test retest reliability of graph metrics in functional brain networks: *A resting-state fNIRS study*. PLoS One, 8(9), e72425. 10.1371/journal.pone.0072425

Öztekin, I., Curtis, C. E., & McElree, B. (2009). The medial temporal lobe and the left inferior prefrontal cortex jointly support interference resolution in verbal working memory. Journal of cognitive neuroscience, 21(10), 1967–1979. 10.1162/jocn.2008.21146

Park, J., & Brannon, E. M. (2014). Improving arithmetic performance with number sense training: An investigation of underlying mechanism. Cognition, 133(1), 188–200. 10.1016/j.cognition.2014.06.011

Pelicioni, P. H., Tijsma, M., Lord, S. R., & Menant, J. (2019). Prefrontal cortical activation measured by fNIRS during walking: effects of age, disease and secondary task. PeerJ, 7, e6833. 10.7717/peerj.6833

Peters, L., Bulthé, J., Daniels, N., de Beeck, H. O., & De Smedt, B. (2018). Dyscalculia and dyslexia: Different behavioral, yet similar brain activity profiles during arithmetic. NeuroImage: Clinical, 18, 663–674. 10.1016/j.nicl.2018.03.003

Pica, P., Lemer, C., Izard, V., & Dehaene, S. (2004). Exact and approximate arithmetic in an Amazonian indigene group. Science, 306(5695), 499–503. 10.1126/science.1102085

Poldrack, R. A., Wagner, A. D., Prull, M. W., Desmond, J. E., Glover, G. H., & Gabrieli, J. D. (1999). Functional specialization for semantic and phonological processing in the left inferior prefrontal cortex. Neuroimage, 10(1), 15–35. 10.1006/nimg.1999.0441

Price, G. R., Yeo, D. J., Wilkey, E. D., & Cutting, L. E. (2018). Prospective relations between resting-state connectivity of parietal subdivisions and arithmetic competence. Developmental Cognitive Neuroscience, 30, 280–290. 10.1016/j.dcn.2017.02.006

Qi, L., Yin, Y., Bu, L., Tang, Z., Tang, L., & Dong, G. (2021). Acute VR competitive cycling exercise enhanced cortical activations and brain functional network efficiency in MA-dependent individuals. Neuroscience Letters, 757, 135969. 10.1016/j.neulet.2021.135969

Rapin, I. (2016). Dyscalculia and the calculating brain. Pediatric neurology, 61, 11–20. 10.1016/j.pediatrneurol.2016.02.007

Räsänen, P., Daland, E., Dalvang, T., Engström, A., Korhonen, J., Kristinsdóttir, J. V., & Träff, U. (2019). Mathematical learning and its difficulties: the case of Nordic countries. International handbook of mathematical learning difficulties: From the laboratory to the classroom, 107-125. 10.1007/978-3-319-97148-3_8

Rivera, S.M., Reiss, A.L., Eckert, M.A., Menon, V., 2005. Developmental changes in mental arithmetic: evidence for increased functional specialization of the left inferior parietal cortex. Cereb. Cerebral cortex, 15(11), 1779–1790. 10.1093/cercor/bhi055

Rubinsten, O., & Henik, A. (2009). Developmental dyscalculia: Heterogeneity might not mean different mechanisms. Trends in cognitive sciences, 13(2), 92–99. 10.1016/j.tics.2008.11.002

Schel, M. A., & Klingberg, T. (2017). Specialization of the right intraparietal sulcus for processing mathematics during development. Cerebral Cortex, 27(9), 4436–4446. 10.1093/cercor/bhw246

Scholkmann, F., Spichtig, S., Muehlemann, T., & Wolf, M. (2010). How to detect and reduce movement artifacts in near-infrared imaging using moving standard deviation and spline interpolation. Physiological measurement, 31(5), 649. DOI 10.1088/0967-3334/31/5/004

Smaczny, S., Sperber, C., Jung, S., Moeller, K., Karnath, H. O., & Klein, E. (2021). Left angular gyrus disconnection impairs multiplication fact retrieval. BioRxiv, 2021–10.

Sokolowski, H. M., Matejko, A. A., & Ansari, D. (2023). The role of the angular gyrus in arithmetic processing: A literature review. Brain Structure and Function, 228(1), 293–304. 10.1007/s00429-022-02594-8

Soltanlou, M., Artemenko, C., Ehlis, A.-C., Huber, S., Fallgatter, A. J., Dresler, T., & Nuerk, H.-C. (2018). Reduction but no shift in brain activation after arithmetic learning in children: A simultaneous fNIRS-EEG study. Scientific Reports, 8(1), 1–15. 10.1038/s41598-018-20007-x

Song, S., & Su, M. (2022). The Intelligence Quotient-math achievement link: evidence from behavioral and biological research. Current Opinion in Behavioral Sciences, 46, 101160. 10.1016/j.cobeha.2022.101160

Storey, J. D. (2002). A direct approach to false discovery rates. Journal of the Royal Statistical Society Series B: Statistical Methodology, 64(3), 479–498. 10.1111/1467-9868.00346

Tak, S., & Ye, J. C. (2014). Statistical analysis of fNIRS data: A comprehensive review. Neuroimage, 85, 72–91. 10.1016/j.neuroimage.2013.06.016

Träff, U., Olsson, L., Östergren, R., & Skagerlund, K. (2017). Heterogeneity of developmental dyscalculia: Cases with different deficit profiles. Frontiers in psychology, 7, 2000. 10.3389/fpsyg.2016.02000

Tressoldi, P. E., Rosati, M., & Lucangeli, D. (2007). Patterns of developmental dyscalculia with or without dyslexia. Neurocase, 13(4), 217–225. 10.1080/13554790701533746

Van den Heuvel, M. P., & Sporns, O. (2013). Network hubs in the human brain. Trends in Cognitive Sciences, 17(12), 683–696. 10.1016/j.tics.2013.09.012

Van Hedel, H. J., Bulloni, A., & Gut, A. (2021). Prefrontal Cortex and Supplementary Motor Area Activation During Robot-Assisted Weight-Supported Over-Ground Walking in Young Neurological Patients: A Pilot fNIRS Study. Frontiers in Rehabilitation Sciences, 93. 10.3389/fresc.2021.788087\

Van Viersen, S., Slot, E. M., Kroesbergen, E. H., Van’t Noordende, J. E., & Leseman, P. P. (2013). The added value of eye-tracking in diagnosing dyscalculia: A case study. Frontiers in psychology, 4, 679. 10.3389/fpsyg.2013.00679

Venkatraman, V., Ansari, D., & Chee, M. W. (2005). Neural correlates of symbolic and non-symbolic arithmetic. Neuropsychologia, 43(5), 744–753. 10.1016/j.neuropsychologia.2004.08.005

Wang, J., Dong, Q., & Niu, H. (2017). The minimum resting-state fNIRS imaging duration for accurate and stable mapping of brain connectivity network in children. Scientific reports, 7(1), 1–10. 10.1038/s41598-017-06340-7

Wang, M., Hu, Z., Liu, L., Li, H., Qian, Q., & Niu, H. (2020). Disrupted functional brain connectivity networks in children with attention-deficit/hyperactivity disorder: evidence from resting-state functional near-infrared spectroscopy. Neurophotonics, 7(1), 015012–015012. 10.1117/1.NPh.7.1.015012

Wang, M. Y., Zhang, J., Lu, F. M., Xiang, Y. T., & Yuan, Z. (2018). Neuroticism and conscientiousness respectively positively and negatively correlated with the network characteristic path length in dorsal lateral prefrontal cortex: A resting-state fNIRS study. Brain and behavior, 8(9), e01074. 10.1002/brb3.1074

Wang, M., Yuan, Z., & Niu, H. (2019). Reliability evaluation on weighted graph metrics of fNIRS brain networks. Quantitative Imaging in Medicine and Surgery, 9(5), 832.

Wechsler, D. J. S. A. P. C. (2014). Wechsler intelligence scale for children–Fifth Edition (WISC-V). Bloomington, MN: Pearson.

Wechsler, D., (2009). Wechsler Individual Achievement Test 3rd Edition (WIAT III). The Psychological Corp, London.

Wong, B. T. M. (2017). Learning analytics in higher education: an analysis of case studies. Asian Association of Open Universities Journal, 12(1), 21–40. 10.1108/AAOUJ-01-2017-0009

Wong, T. T. Y., & Chan, W. W. L. (2019). Identifying children with persistent low math achievement: The role of number-magnitude mapping and symbolic numerical processing. Learning and Instruction, 60, 29–40. 10.1016/j.learninstruc.2018.11.006

Xu, J., Liu, X., Zhang, J., Li, Z., Wang, X., Fang, F., & Niu, H. (2015). FC-NIRS: a functional connectivity analysis tool for near-infrared spectroscopy data. BioMed Research International. 2015(1), 248724. 10.1155/2015/248724.

Zhang, J. X., Zhuang, J., Ma, L., Yu, W., Peng, D., Ding, G., & Weng, X. (2004). Semantic processing of Chinese in left inferior prefrontal cortex studied with reversible words. NeuroImage, 23(3), 975–982. 10.1016/j.neuroimage.2004.07.008

Zhao, H., Li, X., Karolis, V., Feng, Y., Niu, H., & Butterworth, B. (2019). Arithmetic learning modifies the functional connectivity of the fronto-parietal network. Cortex, 111, 51–62. 10.1016/j.cortex.2018.07.016

