## supplementary for "Investigating Frontoparietal Networks and Activation in Children with Mathematics Learning Difficulties: Cases with Different Deficit Profiles"

**Supplementary Information**

**Graph Theory Metrics’ Explanations**

Graph theory (Bollobás, 1985; Bollobás Béla, 2012; Braun et al., 2018), stemming from mathematics field, provides an insight into analysing the brain network, which represents the brain network as set of nodes -- brain regions -- and edges -- pairwise interactions across regions to reveal architectural organizations of the nervous network in the brain (Aarabi & Huppert, 2019). Figure S1 shows an example of a network with eight nodes and ten edges (Van Den & Pol, 2011). Generally, there are two sets of graphical measures for the brain networks: (1) nodal metrics including nodal degree (Nd) and nodal efficiency (NE); (2) global metrics: characteristic path length (Lp) and its normalized characteristic path length (Lambda), network clustering coefficient (Cp) and its normalized clustering coefficient (Gamma), global efficiency (Eglob), local efficiency (Eloc). The definitions and descriptions of these metrics are detailed in Table S1.

**Figure S1**

**Example of a Network**

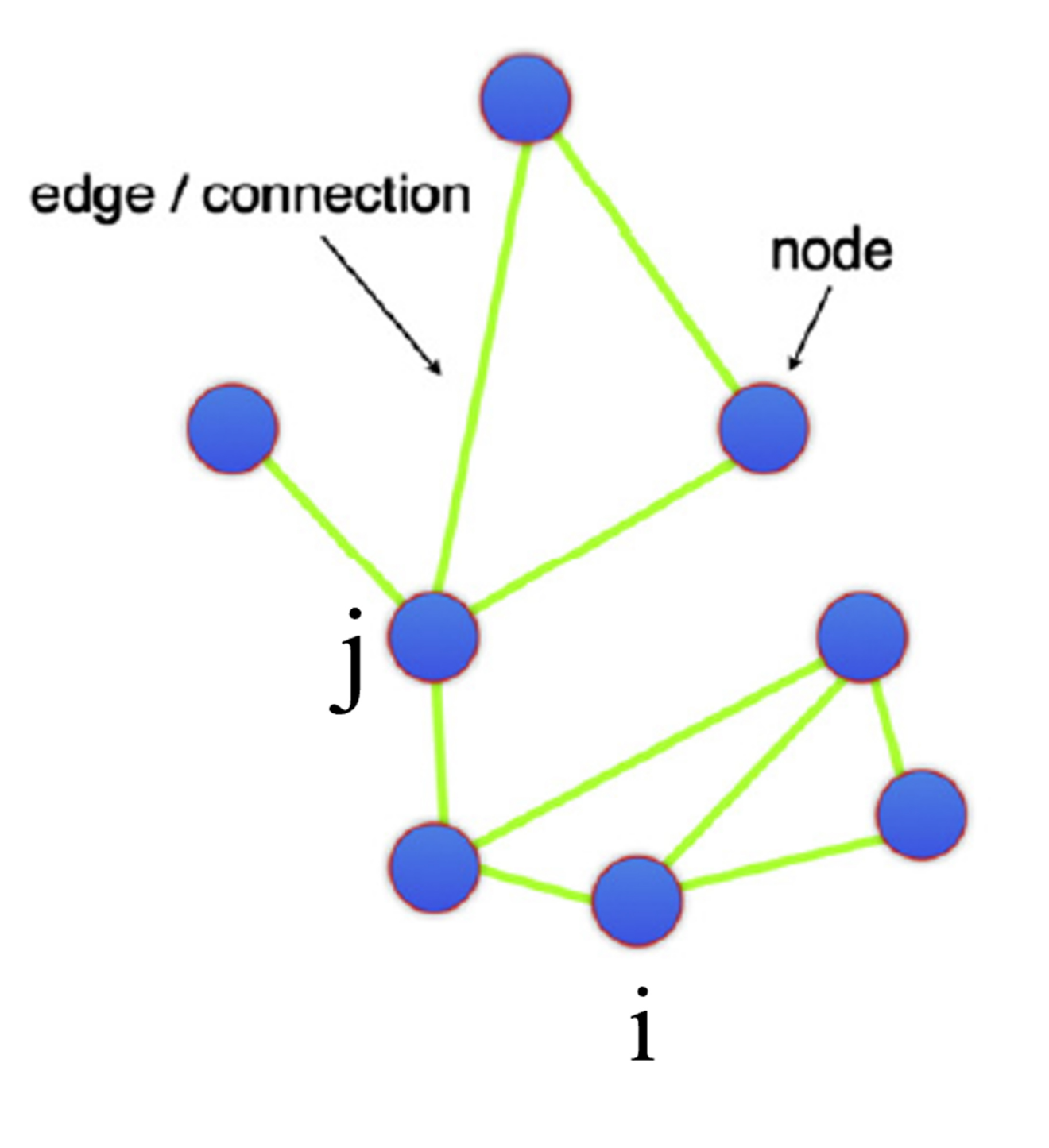

**Table S1**

**Definitions of Graph Theory Metrics**

| Metrics | Definition |
| --- | --- |
| Nodal degree | The number of edges connected directly to a particular node, measures a node’s centrality in a network |
| Clustering Coefficient | The ratio of the total number of edges directedly interconnected among neighbours of a node and the maximum number of directly connected edges among these neighbours. |
| Characteristic Path Length | The shortest path length which denotes the minimum number of path or edges between two nodes. |
| Nodal efficiency | The inverse of the mean of the shortest path length in this network |
| Local efficiency | The inverse of the mean of the shortest path length connecting all neighbours of a node in this subnetwork |
| Global efficiency | The inverse of the mean of the shortest path length of all nodes across the entire brain. |

Nodal Degree (Nd)

Nodal degree, calculated as the number of edges connected directly to a particular node, measures a node’s centrality in a network (Pi et al., 2019). Nodal i in Figure S1 has three degrees. A nodal with higher degree plays a more important role (Aarabi & Huppert, 2019).

Clustering Coefficient (Cp)

Clustering coefficient measures the degree of collectivization in a network and presents the average likelihood that the neighbours of a node were interconnected (Liu, et al, 2023; Pi, et al, 2019). It was computed as the ratio of the total number of edges directedly interconnected among neighbours of a node and the maximum number of directly connected edges among these neighbours. For example, in Figure S1, the node i has three neighbour nodes and there are two directly connected edges among these three nodes. The equation to calculate the clustering coefficient of node i Cp = 2/C (3, 2) = 0.67. The value of the clustering coefficient is not less than 0 (any two neighbours of a node are not directedly connected with each other) and not more than 1 (all neighbours of a node are directedly connected with each other). The clustering coefficient of a network is the average of the cluster coefficients of all nodes in the network.

Characteristic Path Length

Characteristic path length is also termed as the shortest path length which denotes the minimum number of path or edges between two nodes. For instance, the characteristic path length between nodal i and nodal j in Figure S1 is 2. The characteristic path plays an important role in the information transmission of networks. It represents the optimal path serves as a conduit for efficient transmission of neural information across nodes, thereby facilitating energy conservation within the brain system. The characteristic path length of a network is the average shortest path length between every pair of nodes in the network.

Nodal efficiency (NE)

Nodal efficiency quantifies the capability of a particular node transfer information with all other nodes in a network and it is computed as the inverse of the mean of the shortest path length in this network (Geng et al., 2017; Li et al., 2018; Niu et al., 2013).

Local efficiency (Eloc)

Local efficiency quantifies the capability of a subnetwork composed of neighbours of a node to transfer information and it is computed as the inverse of the mean of the shortest path length connecting all neighbours of the node in this subnetwork. The local efficiency of a network is the average of the local efficiency of all nodes in this network.

Global efficiency (Eglob)

Global efficiency quantifies the capability of a network to transfer information globally. It is computed as the inverse of the mean of the shortest path length of all nodes across the entire brain.

**Data exclusion criteria:**

(a) failed to complete the whole procedure; (b) accuracy in addition tasks was less than 3 standard deviations from the mean accuracy of all participants (Moore et al., 2012); (c) the signal-to-noise ratio of resting state was less than 2 (Xu et al., 2015); (d) standard deviation values of neural data during tasks were larger than five standard deviations away from the mean of the moving standard deviation in individual participants (Huppert et al., 2009). Specifically, a total of six children were excluded on criteria (a); one child was excluded on criteria (b); one child was excluded on criteria (c); two children were excluded on criteria (d).

### **Measures**

Wechsler Individual Achievement Test, Third Edition (WIAT-III; Wechsler, 2009)

The WIAT-III was designed to evaluate an individual's academic achievement across a spectrum of domains, encompassing but not limited to reading and mathematics performances. The following seven subtests were administered to assess mathematics and reading composites.

Subtests

Maths Problem Solving. Math Problem Solving primarily measures children’s ability in math reasoning, encompassing tasks that commonly correlate to word problems and the practical application of mathematical operations and principles. Participants were required to solve math word problems presented to them, which could involve a series of sequential actions and pertain to concepts such as time, currency, measurement, probability, geometry, or interpreting graphical representations (Caemmerer et al., 2018). Problems include number recognition and comparison, arithmetic, measuring length and everyday applications, such as “which number is more or less?”, “how many shapes in total?”, “how many centimetres long is the pencil?” and “what time is shown on this clock?”.

Numeracy. This sub-test evaluates children’s proficiency in applying basic arithmetic skills and their understanding of numerical concepts, such as the ability to count, distinguish numbers from letters, recognize mathematics symbols including “+” and “-” and operate with numbers such as “5+1+5+2=?”. The subtest typically includes tasks that vary in complexity and require both rote computation and problem-solving skills (Iseman & Naglieri, 2011).

Maths Fluency. Maths fluency measures children’s proficiency in performing mathematical calculations accurately and efficiently (McCrimmon & Climie, 2011). It often includes tasks such as solving arithmetic problems quickly, which reflects foundational mathematical skills. Participants are required to solve as many arithmetic problems as possible within a set time frame.

Reading Comprehension. This subtest evaluates a student's ability to understand and interpret written materials, including advertisements, informational text and fictional stories (Hulme & Snowling, 2011). It assesses skills such as identifying main ideas, making inferences, drawing conclusions, and summarizing information from these materials through responding oral comprehension questions.

Word Reading. This subtest evaluates children’s ability to decode and read decontextualised words with increasing difficulty levels accurately, which assesses foundational reading skills and the children's knowledge of phonetic and orthographic rules (Griffith, 2016).

Pseudoword Decoding. This subtest measures children’s fluency of decoding nonsense words, such as “ik”, “zad”, “glatch” (Lowell et al., 2014). The pseudowords adhere to English spelling patterns but lack meaningful content (Griffith, 2016).

Oral Reading Fluency. Reading fluency measures the speed, accuracy, and expression when children are reading passages, reflecting the automaticity and efficiency of the reading process and indicating children’s ability to decode words and understand text simultaneously (Lai et al., 2014).

Composites

The scores for each subtest were standardised according to the child’s age group and contributes to composite scores of domain-specific achievements, including Mathematics, Maths Fluency, Total Reading and Basic Reading. Figure S2 details the WIAT-III’s composites derived from combinations of subtests to represent academic domains.

Figure S**2**

WIAT-III Subtests and Composites

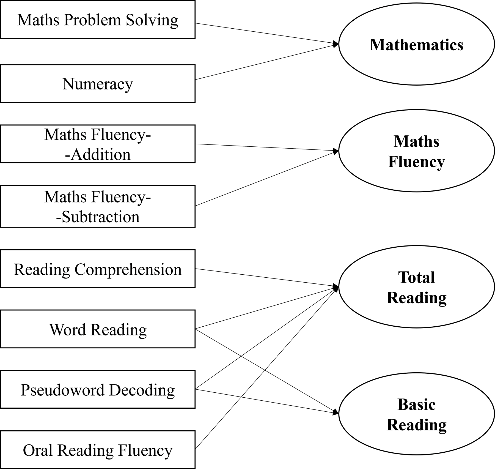

Wechsler Intelligence Scale for Children, **fifth** edition, (WISC-V; Wechsler, **2014**).

The WISC-V is a comprehensive standardized assessment tool designed to measure general cognitive abilities in children between the ages of 6 and 16. It aims to provide a comprehensive evaluation of intellectual potential across a wide range of cognitive domains, including Verbal Comprehension, Working Memory, Processing Speed, Visual Spatial Index, Fluid Reasoning, and Processing Speed through a series of subtests (Gomez et al., 2016).

Ten subtests were administered to assess full scale IQ, including working memory.

Subtests

Block Design. Block Design measures children’s visual-spatial processing and visual-motor coordination (Rader & Hughes, 2005). They were instructed to use either red or white blocks to recreate patterns, which assesses their spatial organization skills and under time constraints.

Similarities. Similarities subtest assesses verbal comprehension and abstract reasoning by requiring children to identify common attributes or conceptual similarities between pairs of words (Díaz et al., 2008). For example, children were required to answer what is the similarity between three and four?

Matrix Reasoning. Matrix Reasoning evaluates children’s pattern recognition, non-verbal reasoning and inductive logic (Chierchia et al., 2019; Weiss, et al., 2013). They were instructed to complete matrices by identifying the missing element through inferring the relationships among various visual shapes.

Digit Span. This subtest assesses children’s working memory and attention span, specifically, their capability of temporarily storing and manipulating auditory information (Lynn & Irwing, 2008). They were instructed to repeat sequences of digits forward and backward and arrange a series of unordered digits in ascending order.

Coding. Coding evaluates children’s processing speed and visual-motor coordination (Oliveras-Rentas et al., 2012). They were instructed to match symbols to geometric shapes including squares, circles and triangles under time constraints.

Vocabulary. This subtest evaluates children’s expressive language and lexical knowledge, including understanding word meanings and linguistic nuances (Wise et al., 2007). They were instructed to explain words presented to them, for example, “what is mouse?”.

Figure Weights. This subtest evaluates children’s quantitative and logical reasoning using contextualized numerical puzzles (Weiss et al., 2016). They were instructed to complete a weight-missing scale by observing the weight relationships between different figures under time constraints.

Visual Puzzles. Visual Puzzles evaluates non-verbal reasoning and spatial visualization by requiring children to reassemble a puzzle that is previously presented to them (Reynolds & Keith, 2017; Weiss et al., 2010). This assesses their skills in perceiving patterns, mental rotation, and deriving solutions within a predetermined time frame.

Picture Span. Picture Span evaluates children’s working memory and attention span in encoding and sequencing visual stimuli, specifically, their capability of temporarily storing and manipulating auditory information (Shapiro, 2017). They were instructed to recall the sequence of presented pictures in the presence of interfering pictures.

Symbol Search. Symbol Search measures children’s visual processing speed and visual discrimination, including rapidly identifying, discriminating, and making decisions about visual stimuli (Keith et al., 2006). They were instructed to identify target symbols in the presence of interfering symbols.

Composites. The score for each subtest was standardised according to children’s age group and contributed to composite scores of domain-general achievements, including Verbal Comprehension, Visual Spatial, Fluid Reasoning, Working Memory, Processing Speed and Full Scale. Figure S3 details the WISC-V’s composites derived from combinations of subtests to represent general cognitive abilities.

Figure S3

WISC-V Subtests and Composites

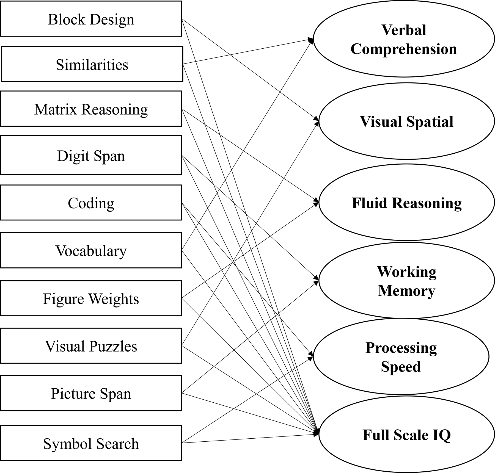

**Figure S4**

**Toys for Freeplay**

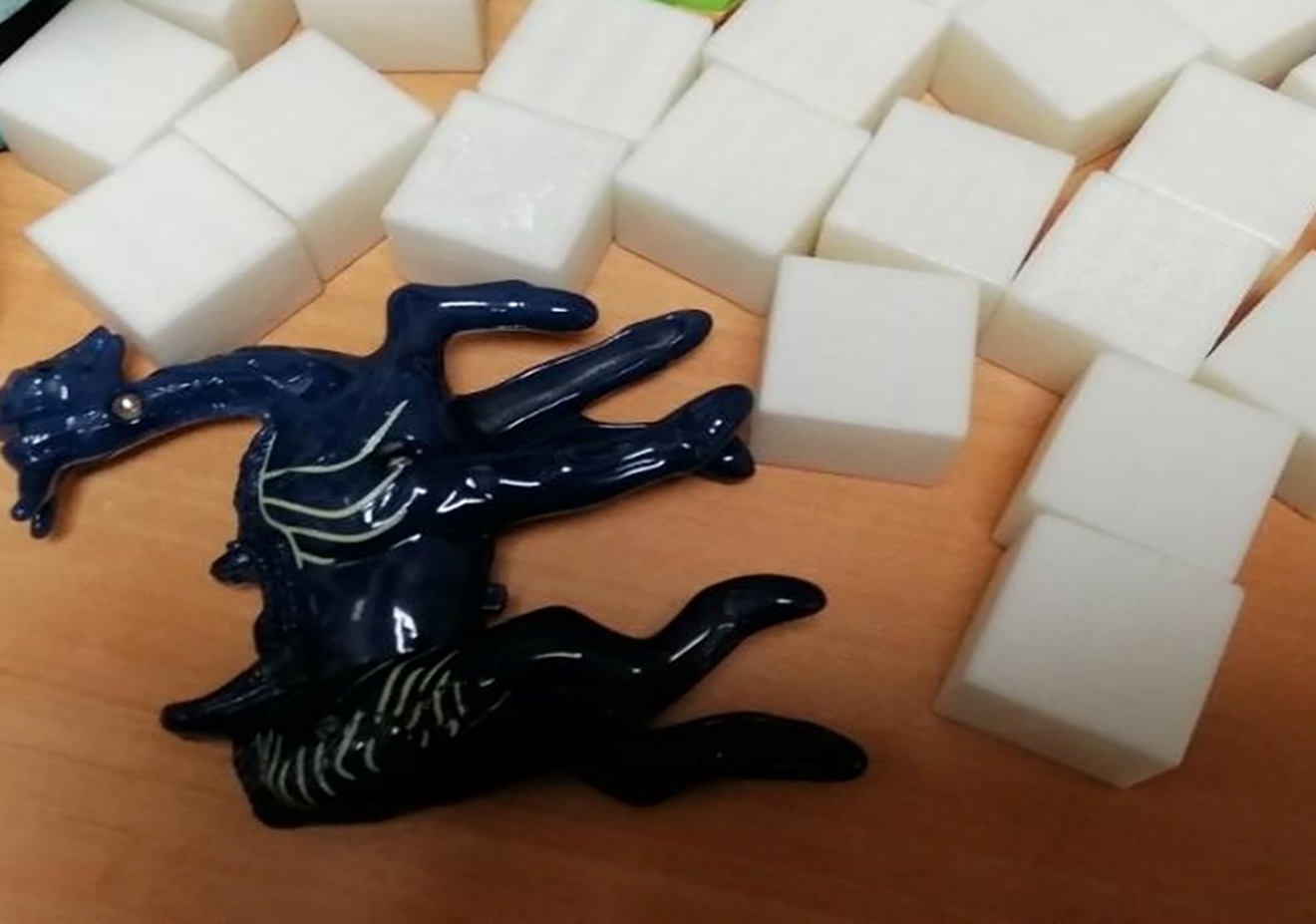

**Table S2**

**Significantly Decreased Activation of Case 2 on the Symbolic Addition Task**

| ID | Brain Area. | HbO conc.  (µmol/L) | TD Mean | TD  SD | T-value | P-value  (Two-tailed) * | Effect Size  (z_cc_) |
| --- | --- | --- | --- | --- | --- | --- | --- |
| Case 2  Boy | LIFG/LMFG (Ch 5) | -0.989 | 0.1326 (43) | 0.2555 | -4.34 | 0.0032 | -4.390 (-5.368 to -3.405) |
|  | LAG (Ch 31) | -0.672 | 0.084  (44) | 0.2173 | -3.440 | 0.0229 | -3.479 (-4.266 to -2.686) |

Note: * Indicated significant p-values after FDR correction.

LIFG: left inferior frontal gyrus; LMFG: left middle frontal gyrus; LAG: left angular gyrus.

**Table S3**

**Global Network Properties of MLD in comparison with TD**

| ID | Global Indicators | TD Mean (N=45) | TD  SD | t-Value | p-value  (Two-tailed) | Effect Size  (z_cc_) |
| --- | --- | --- | --- | --- | --- | --- |
| Case 2  Boy | Gamma (Cp) - 0.8885 | 0.6598 | 0.1017 | 2.224 | 0.0313 | 2.249  (1.693 to 2.797) |
|  | Eloc-0.3151 | 0.2888 | 0.0123 | 2.115 | 0.0401 | 2.138  (1.602 to 2.667) |
|  | Lambda (Lp) - 0.4764 | 0.4322 | 0.0157 | 2.785 | 0.0079 | 2.815  (2.157 to 3.467) |
| Case 3 Girl | Lambda (Lp) -0.4673 |  |  | 2.211 | 0.0323 | 2.236  (1.682 to 2.781) |
|  | Eg - 0.1969 | 0.2243 | 0.01 | -2.710 | 0.0096 | -2.740  (-3.377 to -2.095) |

Note: Eg: global efficiency; Lambda: normalized characteristic path length; Gamma: normalized clustering coefficient; Eloc: local efficiency.

**Table S4**

**Nodal Degree of MLD in comparison with TD**

| ID | Brain Area | Nodal-aND | TD Mean | TD  SD | t-value | p-Value  (Two-tailed) * | Effect Size  (z_cc_) |
| --- | --- | --- | --- | --- | --- | --- | --- |
| Case3  Girl | RMFG (Ch 10) | 0.64 | 6.0950 | 1.5650 | -3.448 | 0.0491 | -3.486  (-4.265 to -2.700) |
| Case 7  Boy | RMFG (Ch 9) | 0.295 | 5.6214 | 1.5052 | -3.5 | 0.0421 | -3.539  (-4.328 to -2.743) |

Note: RMFG: right middle frontal gyrus; aND: Area under curve of nodal degree; ch No.: channel number; * Indicated significant p-values after FDR correction.

**Table S5**

**Nodal Efficiency of MLD in comparison with TD**

| ID | Brain Area | Nodal-aNe | TD Mean | TD SD | t-value | p-value  (Two-tailed) * | Effect Size  (z_cc_) |
| --- | --- | --- | --- | --- | --- | --- | --- |
| Case 2  Boy | RSPG  (Ch 27) | 0.0570 | 0.2314(44) | 0.0365 | -4.725 | 0.0008 | -4.778 (-5.824 to -3.726) |
| Case 3  Girl | LSFG/LMFG  (Ch 1) | 0.0478 | 0.2145(42) | 0.0485 | -3.397 | 0.0149 | -3.437(-4.234 to -2.633) |
|  | SFG/MFG  (Ch 3) | 0.0516 | 0.2380(45) | 0.0357 | -5.164 | 0.0002 | -5.221 (-6.344 to -4.093) |
|  | RMFG  (Ch 10) | 0.0531 | 0.2540(45) | 0.0359 | -5.535 | 0.0002 | -5.596 (-6.794 to -4.392) |
|  | SPG  (Ch 34) | 0.0752 | 0.2040(45) | 0.0419 | -3.04 | 0.0258 | -3.074 (-3.774 to -2.367) |
|  | LSPG  (Ch 35) | 0.0688 | 0.2147(43) | 0.0417 | -3.459 | 0.0149 | -3.499 (-4.298 to -2.692) |
|  | RSPG  (Ch 36) | 0.0855 | 0.2184(43) | 0.0412 | -3.189 | 0.0211 | -3.226 (-3.972 to -2.473) |
| Case 7  Boy | LIFG/LMFG  (Ch 4) | 0.0016 | 0.2269(42) | 0.0562 | -3.962 | 0.0057 | -4.009 (-4.921 to -3.090) |
|  | RMFG  (Ch 9) | 0.0598 | 0.2469(45) | 0.0342 | -5.411 | 0.0004 | -5.471 (-6.644 to -4.292) |
|  | ROG  (Ch 39) | 0.0897 | 0.2228(43) | 0.0424 | -3.103 | 0.0445 | -3.139 (-3.868 to -2.403) |

Note: ROG: right occipital gyrus; RSPG: right superior parietal gyrus; LSFG/LMFG: left superior/middle frontal gyrus; SFG/MFG: superior/middle frontal gyrus; SPG: superior parietal gyrus; LSPG: left superior parietal gyrus; RSPG: right superior parietal gyrus; aNE: Area under curve of nodal efficiency; ch No.: channel number; * Indicated significant p-values after FDR correction.

**References**

Aarabi, A., & Huppert, T. J. (2019). Assessment of the effect of data length on the reliability of resting-state fNIRS connectivity measures and graph metrics. *Biomedical Signal Processing and Control*, 54, 101612. <https://doi.org/10.1016/j.bspc.2019.101612>

Bollobás, B. (1985). Random graphs Academic Press. *New York*.

Bollobás, Béla. (2012). *Graph theory: An introductory course* (Vol. 63). Springer Science & Business Media.

Braun, U., Schaefer, A., Betzel, R. F., Tost, H., Meyer-Lindenberg, A., & Bassett, D. S. (2018). From maps to multi-dimensional network mechanisms of mental disorders. *Neuron*, 97(1), 14–31. <https://doi.org/10.1016/j.neuron.2017.11.007>

Caemmerer, J. M., Maddocks, D. L., Keith, T. Z., & Reynolds, M. R. (2018). Effects of cognitive abilities on child and youth academic achievement: Evidence from the WISC-V and WIAT-III. *Intelligence,* 68, 6-20. <https://doi.org/10.1016/j.intell.2018.02.005>

Chierchia, G., Fuhrmann, D., Knoll, L. J., Pi-Sunyer, B. P., Sakhardande, A. L., & Blakemore, S. J. (2019). The matrix reasoning item bank (MaRs-IB): Novel, open-access abstract reasoning items for adolescents and adults. *Royal Society open science,* 6(10), 190232. <https://doi.org/10.1098/rsos.190232>

Díaz, R., Gual, A., García, M., Arnau, J., Pascual, F., Cañuelo, B., ... & Garbayo, I. (2008). Children of alcoholics in Spain: from risk to pathology: results from the ALFIL program. *Social psychiatry and psychiatric epidemiology,* 43, 1-10. <https://doi.org/10.1007/s00127-007-0264-2>

Geng, S., Liu, X., Biswal, B. B., & Niu, H. (2017). Effect of resting-state fNIRS scanning duration on functional brain connectivity and graph theory metrics of brain network. *Frontiers in Neuroscience*, 11, 392. <https://doi.org/10.3389/fnins.2017.00392>

Gomez, R., Vance, A., & Watson, S. D. (2016). Structure of the Wechsler Intelligence Scale for Children–Fourth Edition in a group of children with ADHD. *Frontiers in Psychology,* 7, 737. <https://doi.org/10.3389/fpsyg.2016.00737>

Griffith, M. E. (2016). *Measuring word reading and pseudoword decoding in struggling readers using the test of word reading efficiency-(TOWRE-2) and the wechsler individual achievement test-(WIAT-III)* (Doctoral dissertation, Middle Tennessee State University).

Hulme, C., & Snowling, M. J. (2011). Children's reading comprehension difficulties: Nature, causes, and treatments. *Current Directions in Psychological Science,* 20(3), 139-142. <https://doi.org/10.1177/0963721411408673>

Huppert, T. J., Diamond, S. G., Franceschini, M. A., & Boas, D. A. (2009). HomER: a review of time-series analysis methods for near-infrared spectroscopy of the brain. *Applied Optics*, 48(10), D280–D298. <https://doi.org/10.1364/AO.48.00D280>

Iseman, J. S., & Naglieri, J. A. (2011). A cognitive strategy instruction to improve math calculation for children with ADHD and LD: A randomized controlled study. *Journal of learning disabilities,* 44(2), 184-195. <https://doi.org/10.1177/0022219410391190>

Keith, T. Z., Fine, J. G., Taub, G. E., Reynolds, M. R., & Kranzler, J. H. (2006). Higher order, multisample, confirmatory factor analysis of the Wechsler Intelligence Scale for Children—Fourth Edition: What does it measure. *School Psychology Review,* 35(1), 108-127.

Lai, S. A., George Benjamin, R., Schwanenflugel, P. J., & Kuhn, M. R. (2014). The longitudinal relationship between reading fluency and reading comprehension skills in second-grade children. *Reading & Writing Quarterly,* 30(2), 116-138. <https://doi.org/10.1080/10573569.2013.789785>

Li, X., Zhu, Z., Zhao, W., Sun, Y., Wen, D., Xie, Y., Liu, X., Niu, H., & Han, Y. (2018). Decreased resting-state brain signal complexity in patients with mild cognitive impairment and Alzheimer’s disease: *A multi-scale entropy analysis. Biomedical Optics Express*, 9(4), 1916. <https://doi.org/10.1364/BOE.9.001916>

Liu, Y., Kang, X. G., Chen, B. B., Song, C. G., Liu, Y., Hao, J. M., ... & Jiang, W. (2023). Detecting residual brain networks in disorders of consciousness: A resting-state fNIRS study. *Brain Research*, 1798, 148162. <https://doi.org/10.1016/j.brainres.2022.148162>

Lowell, S. C., Felton, R. H., & Hook, P. E. (2014). *Basic facts about assessment of dyslexia: Testing for teaching*. Baltimore, Maryland: The International Dyslexia Association, Incorporated.

Lynn, R., & Irwing, P. (2008). Sex differences in mental arithmetic, digit span, and g defined as working memory capacity. *Intelligence,* 36(3), 226-235. <https://doi.org/10.1016/j.intell.2007.06.002>

McCrimmon, A. W., & Climie, E. A. (2011). Test Review: D. Wechsler Wechsler Individual Achievement Test—Third Edition. San Antonio, TX: NCS Pearson, 2009. *Canadian Journal of School Psychology,* 26(2), 148-156. <https://doi.org/10.1177/0829573511406643>

Moore, T., Hennessy, E. M., Myles, J., Johnson, S. J., Draper, E. S., Costeloe, K. L., & Marlow, N. (2012). Neurological and developmental outcome in extremely preterm children born in England in 1995 and 2006: the EPICure studies. *Bmj, 345*.

Niu, H., Li, Z., Liao, X., Wang, J., Zhao, T., Shu, N., Zhao, X., & He, Y. (2013). Test retest reliability of graph metrics in functional brain networks: *A resting-state fNIRS study. PLoS One*, 8(9), e72425. <https://doi.org/10.1371/journal.pone.0072425>

Oliveras-Rentas, R. E., Kenworthy, L., Roberson, R. B., Martin, A., & Wallace, G. L. (2012). WISC-IV profile in high-functioning autism spectrum disorders: impaired processing speed is associated with increased autism communication symptoms and decreased adaptive communication abilities. *Journal of autism and developmental disorders,* 42, 655-664. <https://doi.org/10.1007/s10803-011-1289-7>

Pi, Y.-L., Wu, X.-H., Wang, F.-J., Liu, K., Wu, Y., Zhu, H., & Zhang, J. (2019). Motor skill learning induces brain network plasticity: A diffusion-tensor imaging study. *PloS One*, 14(2), e0210015. <https://doi.org/10.1371/journal.pone.0210015>

Rader, N., & Hughes, E. (2005). The influence of affective state on the performance of a block design task in 6-and 7-year-old children. *Cognition & Emotion,* 19(1), 143-150. <https://doi.org/10.1080/02699930441000049>

Reynolds, M. R., & Keith, T. Z. (2017). Multi-group and hierarchical confirmatory factor analysis of the Wechsler Intelligence Scale for Children—Fifth Edition: What does it measure?. *Intelligence,* 62, 31-47. <https://doi.org/10.1016/j.intell.2017.02.005>

Shapiro, S. M. (2017). Construct validity of the WISC-V picture span subtest (Order No. 10286259). Available from ProQuest Dissertations & Theses Global. (1914674059). Retrieved from <https://remotexs.ntu.edu.sg/user/login?url=https://www.proquest.com/dissertations-theses/construct-validity-wisc-v-picture-span-subtest/docview/1914674059/se-2>

Van Den Heuvel, M. P., & Pol, H. E. H. (2010). Exploring the brain network: a review on resting-state fMRI functional connectivity. *European neuropsychopharmacology,* 20(8), 519-534. <https://doi.org/10.1016/j.euroneuro.2010.03.008>

Xu, J., Liu, X., Zhang, J., Li, Z., Wang, X., Fang, F., & Niu, H. (2015). FC-NIRS: a functional connectivity analysis tool for near-infrared spectroscopy data. *BioMed Research International*. <https://doi.org/10.1155/2015/248724>.

**Wechsler, D. J. S. A. P. C. (2014). *Wechsler intelligence scale for children–Fifth Edition* (WISC-V). Bloomington, MN: Pearson.**

Wechsler, D., (2009). *Wechsler Individual Achievement Test 3rd Edition* (WIAT III). The

Psychological Corp, London.

Weiss, L. C., Holdnack, J. A., Saklofske, D. H., & Prifitera, A. (2016). *Theoretical and clinical foundations of the WISC-V index scores* (pp. 97-121). Amsterdam, The Netherlands: Elsevier Academic Press.

Weiss, L. G., Saklofske, D. H., Coalson, D. L., & Raiford, S. E. (2010). Theoretical, empirical and clinical foundations of the WAIS-IV index scores. In WAIS-IV clinical use and interpretation (pp. 61-94). Academic Press. https://doi.org/10.1016/B978-0-12-375035-8.10003-5

Weiss, L. G., Keith, T. Z., Zhu, J., & Chen, H. (2013). WISC-IV and clinical validation of the four-and five-factor interpretative approaches. *Journal of Psychoeducational Assessment,* 31(2), 114-131. <https://doi.org/10.1177/0734282913478032>

Wise, J. C., Sevcik, R. A., Morris, R. D., Lovett, M. W., & Wolf, M. (2007). The relationship among receptive and expressive vocabulary, listening comprehension, pre-reading skills, word identification skills, and reading comprehension by children with reading disabilities. *Journal of Speech, Language, and Hearing Research*, 50, 1093–1109. <https://doi.org/10.1044/1092-4388(2007/076)>
